# Atrophin-1 Antisense Oligonucleotide Provides Robust Protection from Pathology in a Fully Humanized DRPLA Model

**DOI:** 10.1101/2025.03.18.643143

**Authors:** Velvet L. Smith, Bereket Z. Gidi, Robert M. Bragg, Jeffrey P. Cantle, Aliza Ben- Varon, Briana Nobel, Silvia Prades, Andrea Compton, Julie Greenfield, Joanna Korecka-Roet, Tim Yu, Vikram Khurana, Holly B. Kordasiewicz, Hien T. Zhao, Melissa Barker-Haliski, Daniel D. Child, Jeffrey B. Carroll

## Abstract

Dentatorubral-pallidoluysian atrophy (DRPLA) is a fatal neurodegenerative disease arising from a CAG repeat expansion in the *atrophin-1* (*ATN1*) gene. Because DRPLA, like many repeat expansion disorders (REDs), arises predominantly from toxic gain-of-function mechanisms, we hypothesized that ATN1 knockdown would have therapeutic potential. To test this, we established the first fully-humanized mouse model of a RED, in which one allele of mouse *Atn1* is completely replaced by human *ATN1*, including 112 pure CAG repeats. This novel approach to exploring RED biology provides significant advantages, notably the ability to test sequence-specific therapeutics targeting human sequences, even in introns and untranslated regions of pre-mRNA. We found that our model—the *Atn1^Q112/+^* mouse—recapitulates key features of human DRPLA, including behavioral alterations, reduced brain size and aggregate accumulation. We treated *Atn1^Q112/+^* mice with antisense oligonucleotides (ASOs) targeting mouse *Atn1* (to probe for loss of function concerns), human *ATN1*, or a combination. Treatment with human, but not mouse, *ATN1*-targeting ASOs provides remarkable protection from a range of disease-related behavioral phenotypes, including aggregation of mutant ATN1 (mATN1), and marked rescue of transcriptional dysregulation in the cerebellum. These results have helped motivate an ongoing human clinical study of ASOs targeting *ATN1* for DRPLA.

## Introduction

Dentatorubral-pallidoluysian atrophy (DRPLA) is an invariably fatal neurodegenerative disease caused by the expansion of the CAG tract in the a*trophin-1* (*ATN1*) gene (1, 2). In Japan, DRPLA’s prevalence (0.2–0.7/100,000 people) (3, 4) is comparable with that of Huntington’s disease (HD; 0.4–0.5/100,000) (3, 5, 6), the most common repeat expansion disorder (RED) in North America and Europe (HD prevalence in North America is 8.9/100,000) (3, 4, 7). Conversely, DRPLA is vanishingly rare in Western Europe and North America, with several dozen patients described in the literature and ascertained in an ongoing natural history study (8, 9). The core symptoms of DRPLA are diverse, and include ataxia, cognitive decline, myoclonus, chorea, epilepsy, and psychiatric manifestations (10, 11). Like other REDs, DRPLA shows significant anticipation in the age of onset over generations (1, 10, 12), somatic CAG repeat length instability in the aging brain (13), and intranuclear and cytoplasmic aggregates containing the expanded protein (14). Thus, while DRPLA’s symptoms present a unique challenge compared to other REDs, its underlying mechanisms—and therefore potential approaches to therapy—may be common.

A key challenge in therapeutic development for neurodegenerative diseases is failure of drugs with promising preclinical results to translate into successful human clinical studies (15, 16). Inaccurate disease modeling in mice bears some of the culpability, as mouse models of neurodegeneration often insufficiently replicate key features of disease biology, even with purely genetic forms of neurodegeneration such as DRPLA and HD. Ongoing efforts to improve this situation have seen the steady evolution of mouse models, from transgenic to knock-in mice with humanized mutations in the mouse orthologous gene, which has enabled important disease insights in Alzheimer’s disease (AD) (17, 18), Parkinson’s disease (PD) (19–21), HD (22–24), and several polyglutamine spinocerebellar ataxia’s (SCAs) (25–31). In preclinical studies of AD-associated genetic variants, the need for full genomic contexts has been increasingly recognized, leading to a fully humanized mouse model of *Tau* (32). More recently, the ambitious gene replacement-Alzheimer’s disease (GR-AD) project is generating many humanized mice harboring AD risk genes (33). These mouse models have replaced one of the endogenous murine alleles with the orthologous, full-length human gene, leading to expression of the human gene product in the setting of endogenous murine expression levels and regulatory elements. Full humanization offers clear benefits for oligonucleotide therapeutics, as all potential sequence targets—even those in introns and untranslated regions (UTRs)—can be studied in the mouse context. Furthermore, humanized mice enable targeting phenomena such as alternative splicing and the inclusion of cryptic exons, which often arise from intronic sequences.

Here, we describe the establishment of a novel, fully humanized, DRPLA mouse model (the first fully humanized model of a RED), and the development and preclinical efficacy studies of ATN1-targeting antisense oligonucleotides (ASOs). This work was undertaken as part of a large consortium focused on improving the lives of patients with DRPLA, with several authors (JG, SP, JBC) playing roles as the scientific advisors of CureDRPLA, a non-profit advocacy organization focused on advancing treatments for DRPLA. Since launching in 2019, CureDRPLA has funded an array of tools facilitating work in DRPLA, including Induced Pluripotent Stem Cell Lines, a global natural history study, and an online patient registry (9, 34). These mice, *Atn1^Q112/+^*, are an exciting advancement from existing models of DRPLA (35–37) in that they express the entire human *ATN1* transcript, allowing ASO targeting of any sequence within the gene, from 5’UTR to the 3’UTR. Another advantage is the ATN1 dosage, as all existing DRPLA models rely on transgenic overexpression of some form of mutant ATN1 (mATN1); thus, our model is the first that allows careful analysis of *ATN1* lowering in a genetic context recapitulating individuals with DRPLA.

We hypothesized that reduction of *mATN1* is sufficient to alleviate the DRPLA disease phenotype, and we tested this using ASOs in our humanized DRPLA mouse model. Recently, a distinct clinical syndrome associated with missense mutations in *ATN1* has been described (Congenital hypotonia, epilepsy, developmental delay, and digital anomalies, CHEDDA) (38). While CHEDDA likely arises from a distinct, toxic, *ATN1* gain-of-function (rather than a loss of function), the awareness of two diseases arising from *ATN1* mutations rendered us sensitive to the potential risks of *ATN1* lowering. Thus, we designed ASOs for both human-and mouse-*ATN1* to target each allele individually. We found that treatment with human ATN1 ASO lowers *mATN1* and provides remarkably robust protection from DRPLA-relevant signs, including behavior, neuropathology, and transcriptional dysregulation. Similar rescue is not seen in *Atn1^Q112/+^* mice after silencing WT mouse *Atn1*, nor are disease-relevant phenotypes evidently exacerbated by treatment with the mouse Atn1 ASO. We believe that our results provide a template for other ultra-rare patient advocacy organizations, particularly those focused on neurological conditions amenable to ASO therapy, as these results supported our decision to pursue ASO therapy for DRPLA patients in clinical studies which are underway and will be described separately.

## Results

### Generation of *Atn1^Q112/+^* mice

To screen and test oligonucleotides it is convenient to have a mouse model that expresses the entire target transcript because ASOs can target the entire pre-mRNA (39). To facilitate ASO screening and for maximal construct validity, we generated a mouse in which the endogenous mouse *Atn1* locus was replaced with the human *ATN1* sequence, from 5’UTR to 3’UTR, with expanded (112) sequence-verified pure CAG repeats in exon-5 (Fig. 1). Details of the generation and validation of the *Atn1^Q112/+^* allele are outlined in Fig. S1, and in the materials and methods.

**Figure 1:**
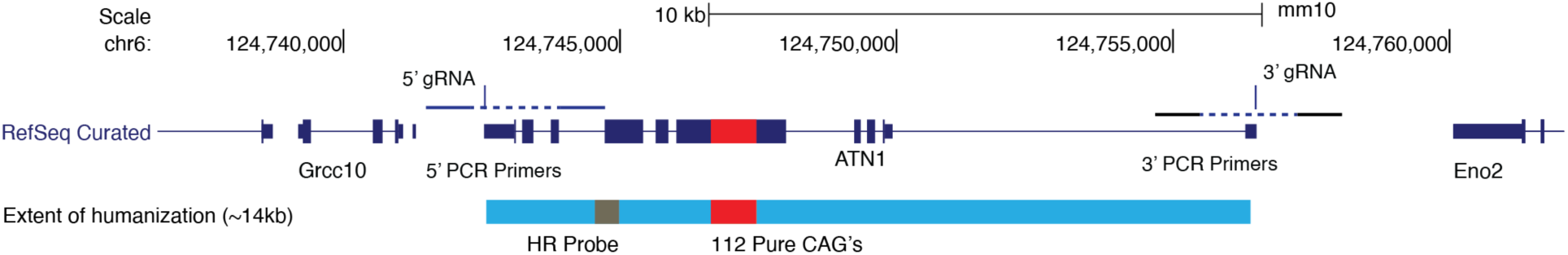
Extent of humanization in the *Atn1^Q112/+^* mouse model of DRPLA, and location of the mouse and human ASO targets. Schematic of the humanized *Atn1* locus at mouse chromosome 6 with the upstream (*Grcc10*) and downstream (*Eno2*) RefSeq curated genes indicated. The 5’ and 3’ gRNA sites used to excise mouse *Atn1* are indicated as labeled vertical dashes, as are PCR primer pairs flanking from the mouse to human sequence at both the 5’ and 3’ ends of the humanized region. Narrow purple rectangles indicate untranslated regions, while thicker bars indicate exons, while introns are indicated by thin lines joining exons. The donor human allele included 112 pure, sequence-verified, CAG repeats, indicated as a red bar. The bright blue bar indicates the extent of humanization, and the relative location of the homologous recombination (HR) probe (dark gray rectangle), and CAG repeats.

### Initial Observations of Emergent Deaths

We generated and shipped an initial cohort of *Atn1^Q112/+^*mice as subjects for an ASO study but found that many mice died in shipment, and the surviving mice were sufficiently ill that they needed to be sacrificed on arrival. This suggests that *Atn1^Q112/+^* mice are sensitive to environmental stress by 8 weeks of age. To proceed with our studies, we conducted *in vitro* fertilization (IVF) with sperm from *Atn1^Q112/+^* founders, resulting in pregnant dams, which were shipped to give birth for pilot studies. We qualitatively observed that phenotypes in these locally born *Atn1^Q112/+^* mice progressed rapidly, including incoordination, gait disturbances, and tremors, particularly in response to sudden noise or movement. The mice were sufficiently uncoordinated in that they struggled to eat and required euthanasia by 10 weeks of age. We attempted to harem breed 6 male *Atn1^Q112/+^* mice with WT females locally with no resulting pups, suggesting that the fertility of *Atn1^Q112/+^* mice is significantly impaired, as has been observed in other transgenic DRPLA mouse models with very long CAG repeats (36). Nevertheless, using IVF, we were able to generate a large cohort of *Atn1^Q112/+^* mice for ASO efficacy studies (Table S1).

### Development of ATN1 Allele Specific ASOs

We developed two ASO compounds—one targeting mouse *Atn1* and one targeting human *ATN1*—by screening ASOs against human or mouse *ATN1* pre-mRNA sequence in cell culture and *in vivo* (using humanized *ATN1* knock-in mice without CAG expansion and WT C57BL/6 mice). The primary outcome for our screen was *ATN1* or *Atn1* transcript levels, as quantified by Quantitative Real-time PCR (qRT-PCR) assays for each screen. Potential lead ASOs were then screened *in vivo* for their acute and chronic tolerability, as well as their potency and selectivity (Figs. S2, S3). We then characterized ASO candidates in a humanized *ATN1* line with non-expanded CAG repeat lengths, enabling us to characterize them *in vivo* without the challenges posed by the severe phenotypes seen in the *Atn1^Q112/+^* mice. Our chosen mouse-and human-*ATN1* ASOs resulted in robust reductions in *Atn1* and *ATN1* levels, respectively, at 2-and 8-weeks post-ICV (intracerebroventricular) injection as determined by allele-selective qRT-PCR (Figs. S2B, S3B). Neither led to any induction of astro-or microglial genes, nor induced any abnormal behaviors when injected in WT or humanized *ATN1* mice after ICV injection of 150 or 300 µg of ASO (Figs. S2C, S3C). We therefore proceeded with a longitudinal study where we tested a mouse *Atn1* targeting ASO, a human *ATN1* targeting ASO, and a combination of both ASOs. We refer to these treatment arms as mASO, hASO, and Combo throughout this manuscript.

### Longitudinal Study Overview

Our human and mouse allele-selective ASOs enabled us to execute a longitudinal characterization of the impact of *ATN1* lowering on emergent behavioral phenotypes in our *Atn1^Q112/+^* mice (Fig. 2A–B). 20 pregnant female mice were generated at Taconic using IVF and shipped to our animal facility at the University of Washington. The dams gave birth approximately five days after their arrival, resulting in a large cohort of well-matched *Atn1^Q112/+^*and WT littermate mice (Tables S2, 3). Thanks to our pilot study observations, we developed a two-stage ASO treatment strategy based on a published approach in a rapidly developing seizure disorder model (40). All mice in our study received an initial ICV injection (2 µL) at postnatal day 1–3 of life of their allotted treatment (saline, mASO, hASO, or Combo) that delivered 30 µg of ASO (the Combo group received 15 µg of each ASO for a total of 30 µg; Fig. 2A–B). At 5 weeks of age, all mice received a second ICV injection (10 µL) containing saline or 200 µg of ASO (the Combo group received 100 µg of each ASO for a total of 200 µg; Fig. 2A– B). We then conducted exhaustive behavioral examination of the mice from 6–8 weeks of age and sacrificed them at approximately 63 days of age (range = 5866). While this study was not powered for a formal survival analysis, we note that of 30 *Atn1^Q112/+^*mice treated with saline or mASO, 10 (33%) died before the end of the study. However, of the 27 *Atn1^Q112/+^* mice treated with hASO (alone or in combination), only 2 died (7%) (Table S4). At sacrifice, we examined human *ATN1* and mouse *Atn1* levels in the entire cohort with allele-selective qRT-PCR and confirmed reduction by 65-and 36% in the cortex, respectively, with no evidence of cross-reactivity of the ASOs (Fig. 2C).

**Figure 2:**
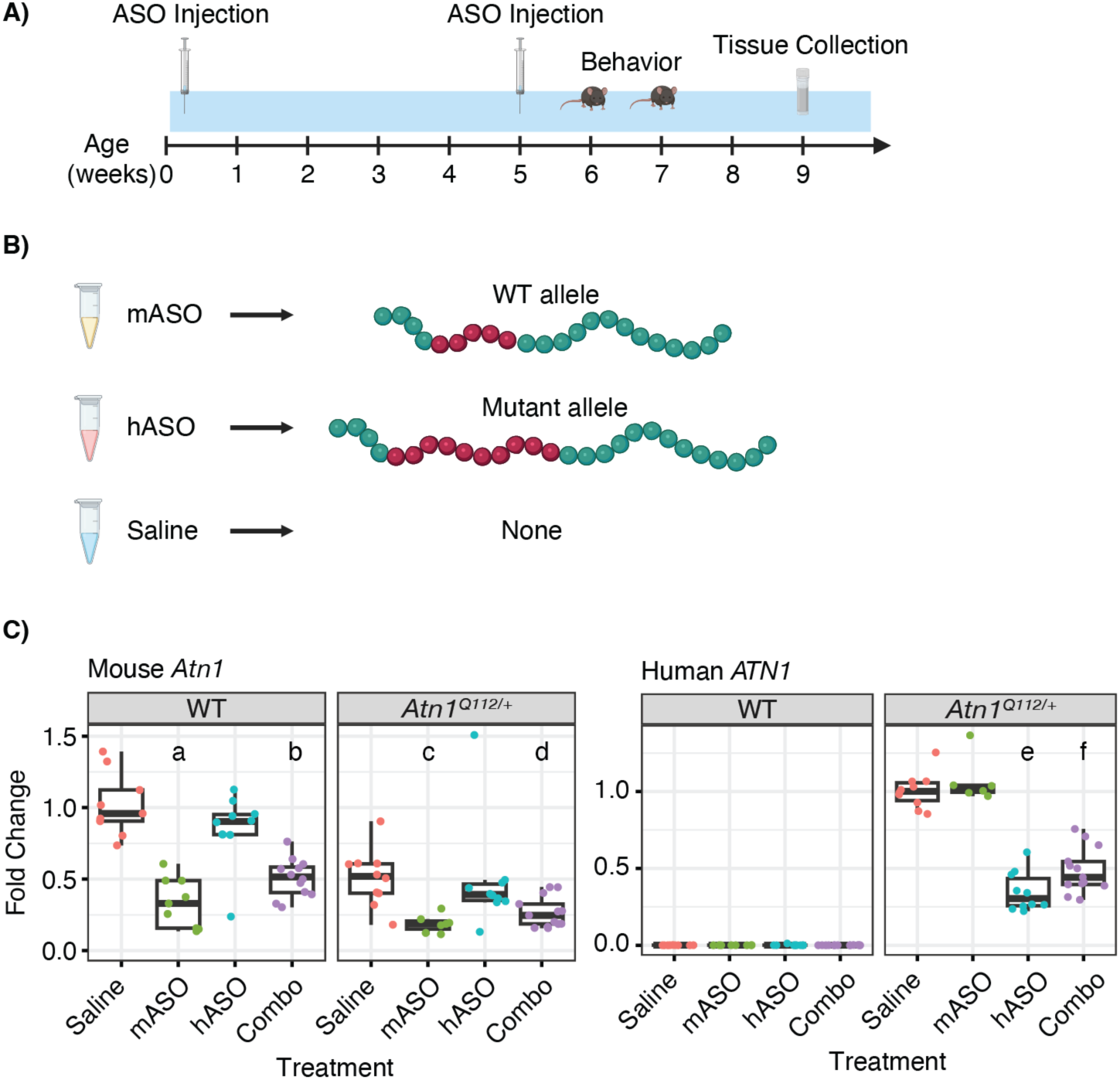
Trial overview and *Atn1/ATN1* lowering achieved in *Atn1^Q112/+^* mice in the study. **A)** Schematic overview of the study design, indicating dates for ASO treatment, behavioral assessments and sacrifice/tissue collection in weeks. **B)** Schematic of the different ASO treatments - the mouse ASO (mASO) only targets mouse *Atn1* and the human ASO (hASO) only targets human *ATN1*. The combination (Combo) of mASO and hASO affects both. **C)** Allele selective mouse and human *Atn1/ATN1* primers were used in qPCR to quantify levels of *Atn1/ATN1* in cortex in all mice from the study outlined in A. In the left plot (n = 103), WT *Atn1* in saline-and mASO-treated *Atn1^Q112/+^* mice is significantly reduced compared to controls (WT mice treatment effect – F_(3,36)_ = 24.8, p = 6.9^-09^; *Atn1^Q112/+^*mice treatment effect – F_(3,36)_ = 4.8, p = 6.6^-03^; select Tukey post-hoc comparison p-values – a - mASO:saline p = 3.8^-08^, b - combo:saline p = 3.9^-06^, c - mASO:saline p = 0.03, d - combo:saline p = 0.06). In the right plot (n = 80), treatment with hASO led to robust reduction in human *ATN1* in *Atn1^Q112/+^* mice when compared to saline-treated *Atn1^Q112/+^*controls (treatment effect –F_(3,35)_ = 66.1, p = 1.7^-14^; select Tukey post-hoc comparison p-values – e - hASO:saline p = 4.0^-12^, f - combo:saline p = 4.0^-10^).

### Human, but not mouse, ASO Rescues Behavioral Abnormalities

We weighed mice weekly throughout the study and detected a complex relationship between sex, genotype, and treatment—in brief, saline-treated *Atn1^Q112/+^* mice were not significantly lighter than sex-matched WT mice (Fig. S4). We conducted a compressed assessment of behavior in the entire cohort of mice from 6–8 weeks of age. Motor coordination assessed with a tapered balance beam task and fixed-speed rotarod task at 4-and 8-rpm, all revealed profound deficits in saline-and mASO-treated *Atn1^Q112/+^* mice (Fig. 3B, 3D). These motor deficits were robustly rescued in *Atn1^Q112/+^* mice treated with hASO. We used an open field arena to measure activity levels and found that saline-treated *Atn1^Q112/+^* mice were significantly hypoactive compared to controls, with a striking number of freezing episodes (205.8% increase in freezing episodes; Figs. 4A, S6D). This hypoactive behavior was significantly ameliorated in mice treated with hASO. We also found that *Atn1^Q112/+^* mice treated with hASO travelled a greater distance during open field testing compared to *Atn1^Q112/+^* mice treated with saline (Figs. 3C, 4B). Next, we conducted a modified SHIRPA (41, 42), which broadly characterize the neurological health of mice (Table S5). This evaluation revealed profound changes in lethargy, tremors, gait, pelvic elevation, and touch escape in saline-and mASO-treated *Atn1^Q112/+^* mice—all of which were markedly rescued by treatment with hASO (Figs. 3A, S5A–F, S6A–C). We saw similar improvements in circadian behavior in a subset of *Atn1^Q112/+^* mice treated with hASO compared to saline examined over 67 hours (Figs. S7, S8). Overall, treatment with the hASO, but not mASO, led to robust protection from deficits in the SHIRPA (Fig. 3A), balance beam (Fig. 3D), rotarod (Fig. 3B), open field assays (Figs. 3C, 4, S6D), and circadian rhythm (Figs. S7, S8).

**Figure 3:**
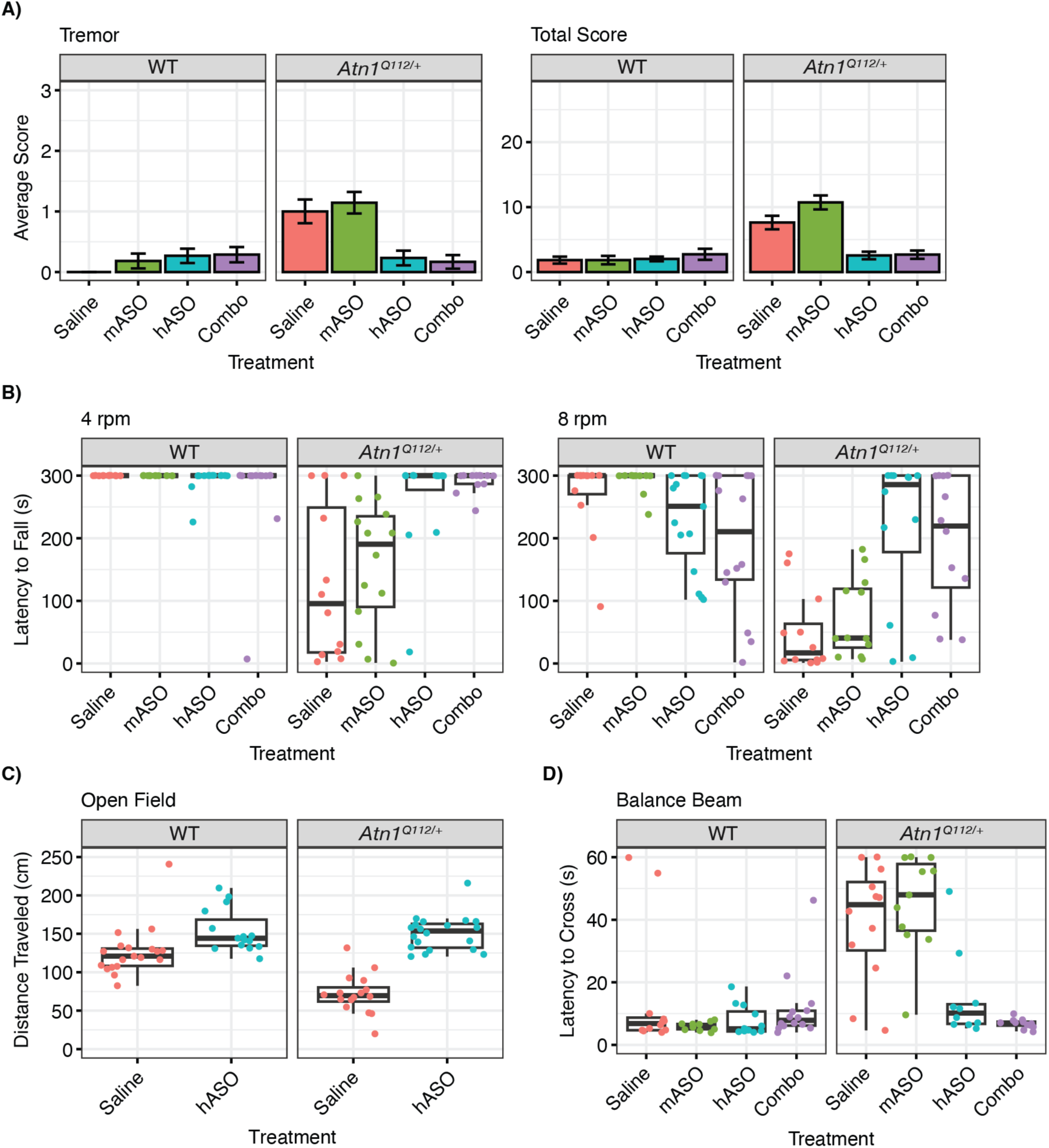
Human, but not mouse, *ATN1* ASO treatment improves behavioral deficits in the *Atn1^Q112/+^*mice. **A)** Bar plots displaying mean score from SHIRPA observational testing of tremors (left) and the total score (right; n = 103, remaining endpoints in Figs. S1–S2C). Total score was calculated by taking the mean of the summed score of all observations with the error bars indicating standard error. Tremor and total score were significantly worsened in mASO and saline-treated *Atn1^Q112/+^* mice when compared to WT mice. Both scores improved with hASO treatment (GLM ANOVA results—genotype effect – *X^2^* (1) = 13.4, p = 2.6^-04^; treatment effect – *X^2^* (3) = 8.2 p = 4.1^-02^; interaction effect – *X^2^* (3) = 14.1, p = 2.7^-03^). **B)** Average time to fall in a fixed rotarod test with the average of three testing trials plotted in seconds at 4-rpm (left, n = 101) and 8-rpm (right, n = 99). *Atn1^Q112/+^*mice treated with mASO and saline fell off the rod significantly earlier than WT mice at both speeds, which is rescued by treatment with hASO, alone or in combination (4-rpm—genotype effect – F_(1,93)_ = 34.9, p = 5.7^08^, treatment effect – F_(3,93)_ = 5.6, p = 0.001, interaction effect – F_(3,93)_ = 9.8, p = 1.12^-05^; 8-rpm—genotype effect – F_(1,91)_ = 39.0, p =1.32^-08^, treatment effect – F_(3,91)_ = 2.1, p = 0.11, interaction effect – F_(3,93)_ = 12.8, p = 4.6^-07^). **C)** Total distance traveled in an open field assessment, measured in centimeters (cm) (n = 70). *Atn1^Q112/+^* mice treated with saline are hypoactive compared to WT, which is reversed by hASO treatment (genotype effect – F_(1,66)_ = 12.2, p = 8.4^-4^, treatment effect – F_(1,66)_ = 66.6, p = 1.4^-11^, interaction effect – F_(1,66)_ = 15.5, p = 2.0^-04^). **D)** Latency to cross tapered balance beam testing with the average of three testing trials plotted in seconds (n = 93). Saline-and mASO-treated *Atn1^Q112/+^*mice are slow to cross an elevated balance beam, which is rescued by treatment with hASO (genotype effect – F_(1,85)_ = 37.3, p = 2.3^-8^, treatment effect – F_(3,85)_ = 11.7, p = 1.7^-6^, interaction effect – F_(3,85)_ = 12.9, p = 4.9^-07^).

**Figure 4:**
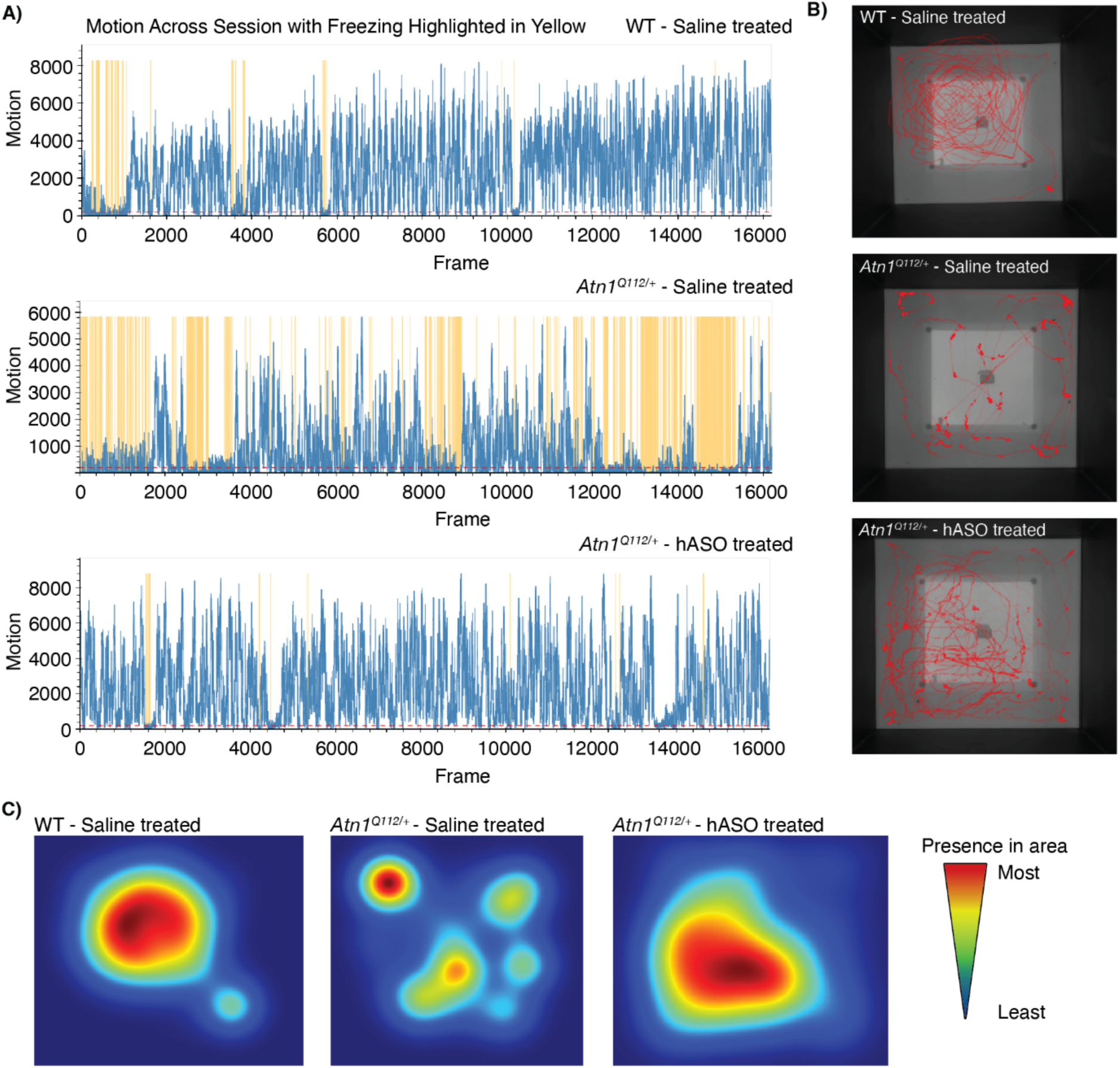
Human *ATN1* ASO treatment rescues locomotor deficits in the *Atn1^Q112/+^*mice. **A)** Representative line graphs of motion over 9 minutes of open field testing (n = 1 per plot). Freezing is indicated in yellow while movement is in blue. The WT saline-treated mouse (top) displayed a high degree of movement along the full-time course, whereas the saline-treated *Atn1^Q112/+^* mouse (middle) had more freezing episodes. The *Atn1^Q112/+^* mouse treated with hASO displayed a very similar freezing pattern to the WT saline-treated mouse. This data is quantified in Fig. S2D for all mice tested. **B)** Representative motion traces from 9 minutes of open field testing (n = 1 per image). The *Atn1^Q112/+^*hASO-treated mouse (bottom) travelled a larger distance than the saline-treated *Atn1^Q112/+^* mouse (middle) and is comparable to the WT saline-treated mouse. This data is quantified in Fig. 3C. **C)** Representative heatmap displaying the presence of mice in 9 minutes of open field testing (n = 1 per image). Mouse presence in a certain area of the box increases as color moves from blue to red. The WT saline-treated mouse (left) was present in many different locations within the box whereas the *Atn1^Q112/+^* saline-treated mouse (middle) spent most of its time in one corner of the box. The *Atn1^Q112/+^* hASO-treated mouse (right) followed a similar pattern to the WT saline-treated mouse.

### Human, but not mouse, ASO Rescues Brain Weight and Aggregate Accumulation

At sacrifice we collected terminal plasma (via cardiac puncture), several peripheral organs, and the brains of each animal in the study. We weighed the whole brain and observed a decrease in saline-treated *Atn1^Q112/+^*mice compared to WT littermates (Fig. 5A), which was not observed when normalized to their body weight (Fig. 5B). Neurofilament light (NEFL) is an axonal protein whose plasma levels are routinely used as a biomarker of ongoing neural damage (45–47).

**Figure 5:**
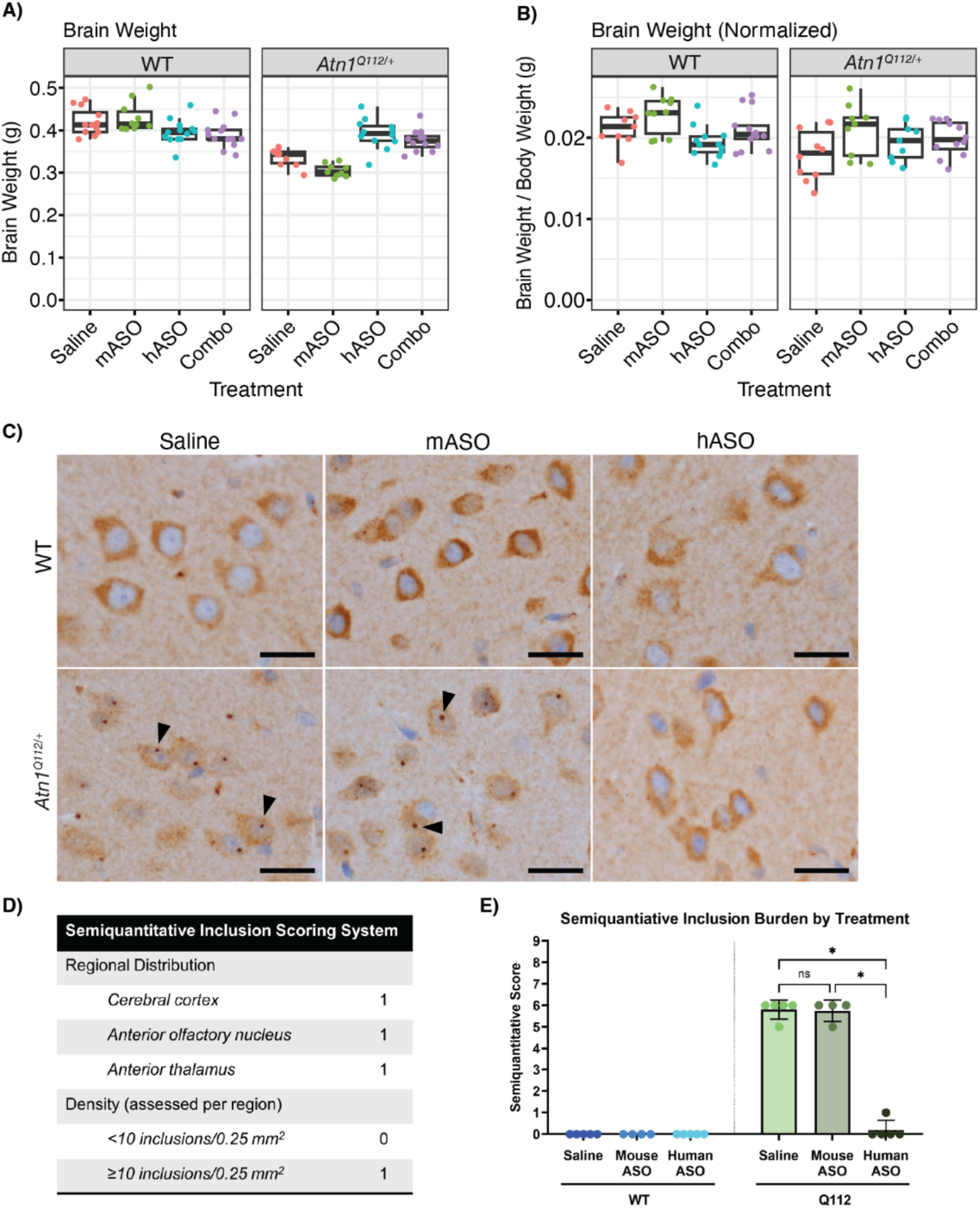
Human, but not mouse, *ATN1* ASO treatment improves gross neuropathological deficits in the *Atn1^Q112/+^*mice. **A)** Raw brain weight (n = 91) is reduced in saline-and mASO-treated *Atn1^Q112/+^* mice, which is rescued by hASO treatment (genotype effect – F_(1,76)_ = 77.5, p = 3.2^-13^, treatment effect – F_(3,76)_ = 1.0, p = 0.4, interaction effect – F_(3,76)_ = 20.3, p = 9.5^-10^). **B)** When normalized to body weight, more subtle impacts of treatment and genotype remain (n = 84; genotype effect – F_(1,76)_ = 7.9, p = 6.2^-3^, treatment effect – F_(3,76)_ = 4.4, p = 6.7^-3^, interaction effect – F_(3,76)_ = 1.5, p = 0.22). **C)** Representative light microscopy images of sections stained with p62 antibody, highlighting the presence or absence of cortical neurons inclusions in each treatment group. Scale bars = 20 µm. **D)** Semiquantitative scoring system designed to assess neuronal inclusion burden in *Atn1^Q112/+^* mouse brains, encompassing distribution and density of affected neurons. **E)** Mean semiquantitative inclusion burden scores across treatment groups. Error bars indicate standard deviation. *p < 0.05, Kruskal-Wallis test followed by Dunn multiple comparison analysis.

Surprisingly, given the severity of their neurological phenotypes, NEFL levels are normal in our *Atn1^Q112/+^* mice, nor did we observe any effect of ASO treatment (Fig. S9). To examine the impact of ASOs on neuropathological phenotypes, we prepared parasagittal brain sections at the level of the mid-olfactory bulb from 4–5 mice per treatment group. These sections were stained with either H&E or by immunohistochemistry for GFAP, IBA1, NeuN, and p62. A board-certified neuropathologist (DDC) blindly examined all sections. Consistent with our NEFL observations, there were no signs of neuron loss noted in *Atn1^Q112/+^* mice, nor changes in neuron density or distribution (Fig. S10A), microvacuolization (Fig. S10B), reactive gliosis (Fig. S11A), or microglial nodules (Fig. S11B). Interestingly, no signs of autophagy adapter p62^+^ aggregates were observed in the regions classically affected in DRPLA, including the globus pallidus, subthalamic nucleus, red nucleus, or deep cerebellar nuclei. However, we observed robust accumulation of p62-immunoreactive inclusions in neurons throughout the cerebral cortex (involving the entire motor, sensory, and visual regions), anterior olfactory nucleus, and anterior thalamus. We observed peri-nuclear and nuclear puncta only in the brains of saline-treated *Atn1^Q112/+^* mice, the severity of which are not impacted by mASO treatment (Fig. 5C); however, we observed minimal inclusions in hASO-treated brains (Fig. 5C). Based on the appearance of our tissue, we devised a semi-quantitative scoring system that incorporated both regional distribution and inclusion burden (Fig. 5D). Scoring each brain by this system demonstrated similar inclusion burden within the saline and mASO-treated *Atn1^Q112/+^* mouse brains, but nearly complete resolution of inclusions in *Atn1^Q112/+^* mouse brains treated with hASO, which were similar in appearance to WT mice (Fig. 5E).

### Human, but not mouse, ASO Rescues Transcriptional Features

While ATN1’s precise roles remain somewhat obscure, it is widely accepted to act as a transcriptional regulator (48). Consequently, we conducted bulk RNA sequencing (RNASeq) of cerebellar tissue from a subset of mice in this study (total *N* = 30; 5 per arm, excluding Combo). In saline-treated *Atn1^Q112/+^* animals, we observe widespread transcriptional dysregulation, with a total of 5,423 genes showing evidence of dysregulation at a permissive threshold of FDR < 0.1 compared to WT; while 1,041 transcripts are downregulated, 655 upregulated at a more conservative fold change |Log_2_FC| > 0.58 and FDR < 0.05 threshold (Fig. 6A). Unsupervised clustering of differentially expressed genes (DEGs) cleanly separated the genotypes (Fig. 6B). Over-representation analysis using `enrichR` (49–51) revealed that DEGs in the *Atn1^Q112/+^* cerebellum are enriched for functional pathways related to neuronal function, including KEGG pathways for calcium signaling (27/189 overlap, p = 0.005), glutamatergic synapse (18/114 overlap, p = 0.007), and axon guidance (25/80 overlap, p = 0.009; Fig. 6C). Of particular interest, given roles for ATN1 in regulating gene expression, DEGs are enriched for PRC2 target genes, with enrichment for genes occupied by SUZ12 (244/1684 overlap with SUZ12 ChEA, p = 4.3^-18^) and EZH2 (34/237 overlap with EZH2 ChEA, p = 0.002; Fig. S12C). We also observe robust enrichment in gene expression studies from HD, suggesting some common transcriptional impacts of CAG expansion (Figs. S12, S13) Turning to the impact of ASO treatment, we first noticed that hASO treatment reduced the number of downregulated DEGs at our stringent threshold nearly in half (i.e. from 1,041 in saline-treated *Atn1^Q112/+^* to 527 in hASO-treated *Atn1^Q112/+^* compared to saline-treated WT; Fig. 7A), with more modest rescue seen in upregulated DEG numbers (from 655 to 529) and only small changes in DEGs of mASO-treated mice. Overlap analysis determined a core set of dysregulated genes in *Atn1^Q112/+^* mice that did not respond to treatment, as well as a large set of dysregulated genes shared by saline-treated and mASO-treated *Atn1^Q112/+^* mice, but not hASO-treated (Fig. 7B). Examples of genes with expression profiles rescued to near WT levels in *Atn1^Q112/+^* mice following hASO treatment include neuronal signaling genes such as *Penk* and *Gpr63* (Fig. 8A). Unsupervised complete linkage clustering of cerebellar DEGs grouped samples primarily by genotype, though we observe treatment-level effects as well (Fig. 7C).

**Figure 6:**
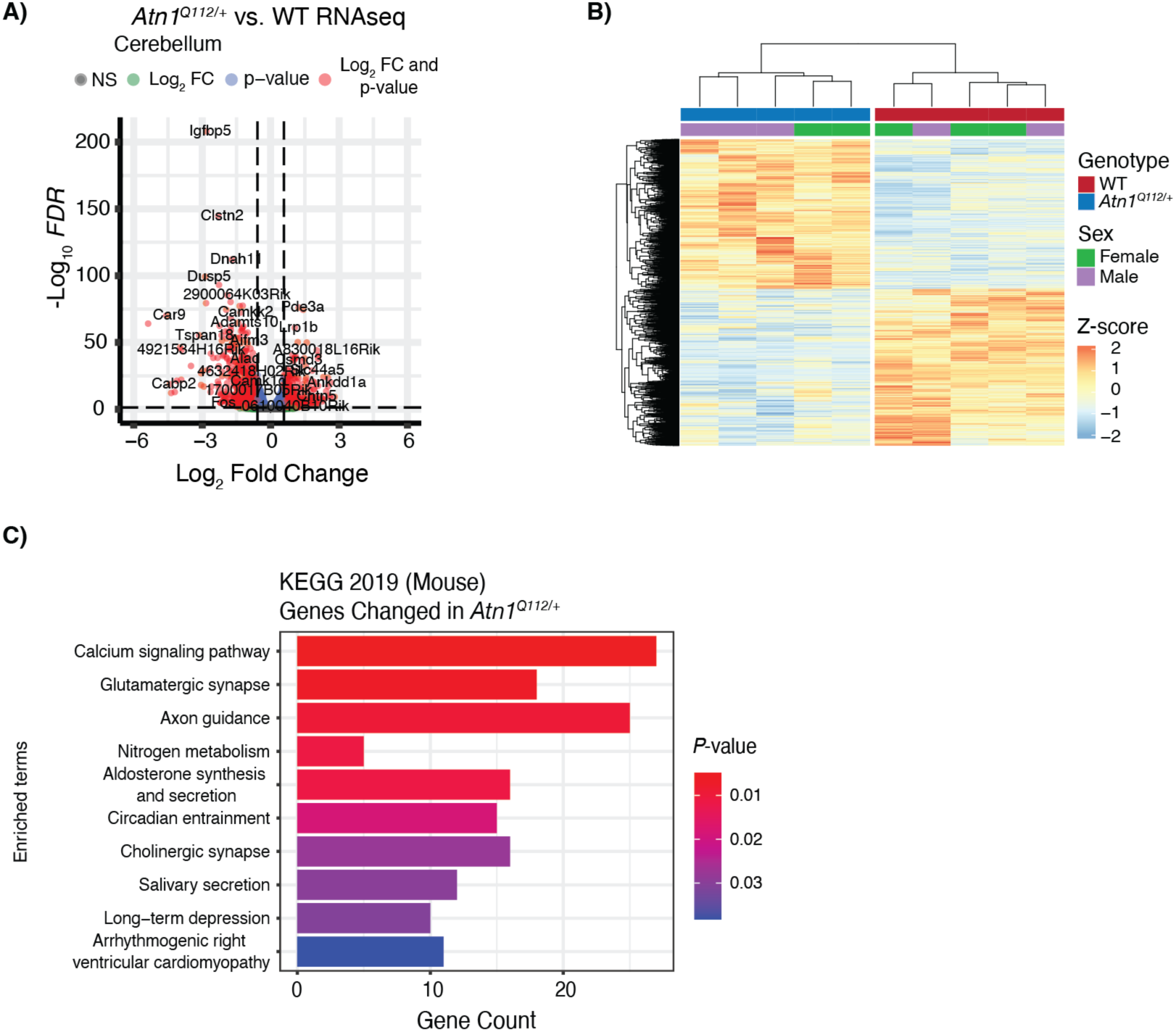
Saline-treated *Atn1^Q112/+^* mice exhibit robust cerebellar transcriptional dysregulation. **A)** A volcano plot showing the severity of transcriptional dysregulation in the cerebellum of saline-treated *Atn1^Q112/+^*mice (n = 5) compared to saline-treated WT mice (n = 5), as quantified with bulk RNA sequencing. Dashed horizonal line indicates an FDR corrected p-value of 0.05, while vertical dashed lines indicate log_2_ fold change of +/-0.58, or roughly 1.5-fold dysregulation. A majority of transcriptionally dysregulated genes in *Atn1^Q112/+^* mice are downregulated. **B)** Unsupervised clustering of the 5,413 DEGs reaching the threshold of FDR < 0.1 in saline-treated *Atn1^Q112/+^*mice compared to saline-treated WT. Each column represents a mouse, while every row indicates a gene. Mice clustered by genotype, indicating that the *Atn1^Q112/+^*transcriptional profile is robust and reproducible. **C)** Geneset over-representation analysis on cerebellar DEGs, finds enrichment for KEGG pathways related to neuronal function such as calcium signaling, glutamatergic synapse, and axon guidance.

**Figure 7:**
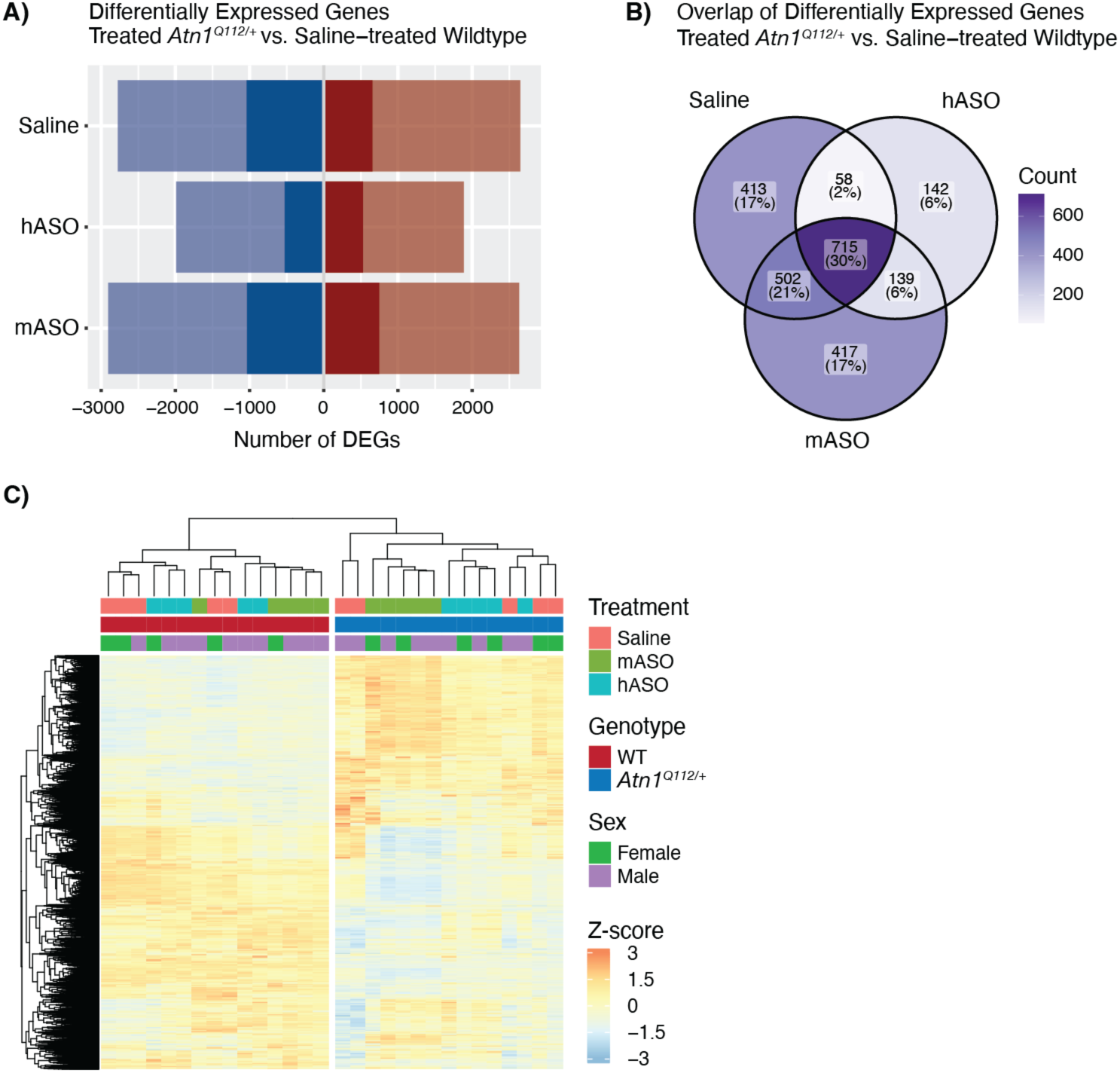
Treatment with human *ATN1* ASO leads to normalization of transcriptional deficits in differentially expressed genes. **A)** Counts of cerebellar down-(blue) and up-regulated (red) DEGs in saline-or ASO-treated *Atn1^Q112/+^* mice (n = 5 for each treatment) compared to saline-treated WT (n = 5) mice. Lighter shading indicates the DEG count with a more permissive threshold of FDR < 0.1, while darker shading indicates a more stringent FDR < 0.05 and |LFC| > 0.58. Most DEGs are downregulated with a significant difference in the number of DEGs in *Atn1^Q112/+^* mice treated with hASO. **B)** Venn diagram indicating DEGs in *Atn1^Q112/+^* mice with the indicated treatment group compared to saline-treated WT. Darker colors indicate more overlap. **C)** Heatmap with unsupervised clustering shows grouping by genotype.

**Figure 8:**
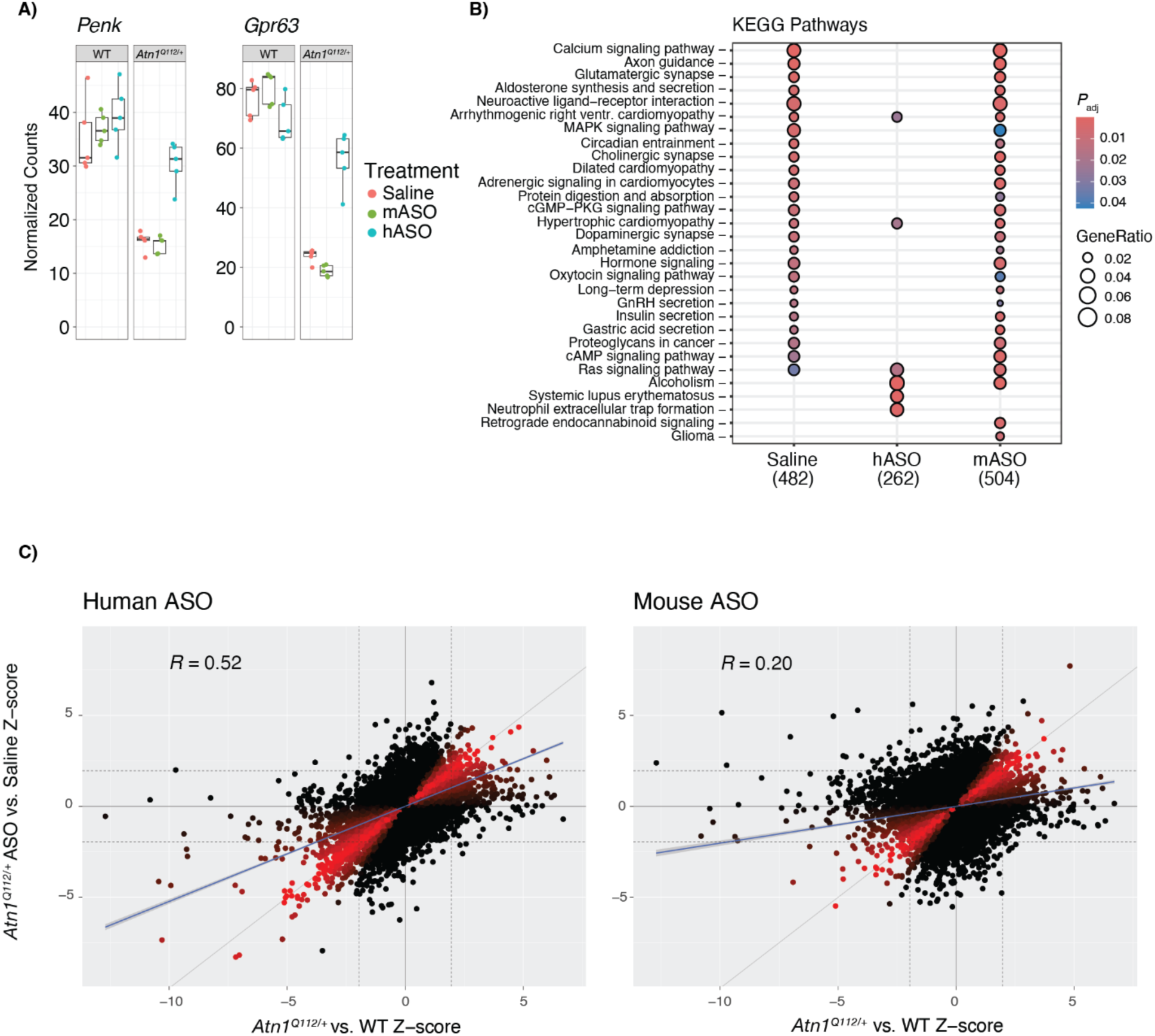
Treatment with human *ATN1* ASO leads to normalization in transcriptional deficits in important genes and pathways in *Atn1^Q112/+^* mice. **A)** Individual gene plots of cerebellar RNA sequencing counts from proenkephalin (*Penk*) and G protein-coupled receptor 63 (*Gpr63*). Saline-or mASO-treated *Atn1^Q112/+^*mice show reduced counts for both genes when compared to WT saline-treated (*Penk*: *Atn1^Q112/+^* saline-treated versus WT saline-treated *padj* = 1.07^-18^, *Atn1^Q112/+^* mASO-treated versus WT saline-treated *padj* = 6.07^-22^; *Gpr63*: *Atn1^Q112/+^*saline-treated versus WT saline-treated *padj* = 8.63^-60^, *Atn1^Q112/+^*mASO-treated versus WT saline-treated *padj* = 1.17^-84^), while *Atn1^Q112/+^* mice treated with hASO have counts significantly restored from *Atn1^Q112/+^* saline-treated (Penk: *Atn1^Q112/+^*hASO-treated versus *Atn1^Q112/+^* saline-treated *padj* = 4.33^-10^; Gpr63: *Atn1^Q112/+^* hASO-treated versus *Atn1^Q112/+^*saline-treated *padj* = 1.16^-28^), and in the case of *Penk* is indistinguishable from WT saline-treated mice (*Atn1^Q112/+^*hASO-treated versus WT saline-treated *padj* = 0.12). **B)** clusterProfiler plot showing enrichment of DEGs in *Atn1^Q112/+^* mice with the indicated treatment group compared to saline-treated WT (FDR < 0.05, |LFC| > 0.58). **C)** Scatterplots comparing scaled cerebellar differential gene expression (z-scores) between *Atn1^Q112/+^* mice and saline-treated WT (x-axis) to z-scores for *Atn1^Q112/+^* mice treated with hASO (left plot) or mASO (right plot) versus saline-treated WT (y-axis each respective plot). Each point represents one gene, and dashed lines indicate z-scores of +/-2.54, encompassing ∼99% of the data range in each dimension. Complete rescue of transcriptional dysregulation would appear as correlation of *R* = 1 (indicated by a gray line with slope = 1), and genes with expression levels returning to WT levels in treated *Atn1^Q112/+^* mice are expected to be closer to this hypothetical line. Linear regression was performed (blue line), with hASO treatment results in *R* = 0.52, while mASO shows less return to WT-like expression, *R* = 0.20. Datapoints with z-score ratio 0 < x:y < 2, have been color heatmapped with decreasing red brightness as values diverge from 1. n = 5 per condition for all panels.

Using the more stringent FDR and fold change threshold noted above, we visualized enriched KEGG pathways using the `compareCluster` function in `clusterProfiler` (52), finding near complete loss of enrichment of DEGs in critical neuronal function categories such as *calcium signaling pathway*, *axon guidance*, and *glutamatergic synapse* following hASO treatment (Fig. 8B). Plotting the z-score of differential gene expression in saline-treated *Atn1^Q112/+^*mice versus saline WT (x-axis) against ASO-treated *Atn1^Q112/+^*mice versus saline-treated WT (y-axis), highlights that hASO treatment restores a larger number of genes to near-WT levels (hASO *R* = 0.52, Fig. 8C) than mASO treatment (mASO *R* = 0.20; Fig. 8C).

## Discussion

In this study we established a novel fully humanized mouse model of DRPLA, developed ASOs targeting mouse and human *ATN1*, and executed a study of their efficacy in our new model. Our results reveal that expressing human *ATN1* with 112 CAG repeats induces severe DRPLA-relevant phenotypes in mice, and that lowering *mATN1* provides robust protection from these alterations.

Excitement about ASO therapy has grown from the striking rescue provided by ASO treatment to children with spinal muscular atrophy by Nusinersen, a splice modulating ASO (53). Subsequently, an ASO developed for familial Amyotrophic lateral sclerosis (ALS) caused by mutations in the *FUS* gene robustly reduces levels of FUS using a novel gapmer ASO modality, improving ALS severity scores, and markedly reducing NEFL levels in the plasma of a single patient (54). Additional positive data was provided by an ASO targeting SOD1 for use in patients with SOD1-mutation-associated ALS (SOD1-ALS; Tofersen). Tofersen treatment robustly reduces SOD1 levels in cerebral spinal fluid (CSF), nominally improves ALS functional scales, and markedly lowers NEFL levels in SOD1-ALS patients (55, 56). While not achieving significance on its primary clinical endpoint in a phase 3 study, Tofersen was recently approved for SOD1-ALS by the U.S. FDA and European Medicines Agency based on the wealth of supportive data suggesting it may modify the disease course in these patients (57). Less positively, an ASO for HD (Tominersen) that targets the Huntingtin (*HTT*) transcript dose-dependently reduces mutant HTT (mHTT) levels in the CSF of treated patients (58), but led to ventricular enlargement, transient increases in NEFL levels, and exacerbation of HD symptoms, leading to an early halt to its phase 3 study (59).

Given this long experience with ASO drugs for genetic CNS diseases, the FDA has provided guidance to allow investigators interested in treating patients impacted by rare diseases like DRPLA to run “*N*-of-1” trials (60). These *N*-of-1 trials give patients with ultra-rare diseases a path toward treatment. As of now, this approach is the only way forward, in our opinion, to provide treatment for DRPLA patients living in the United States (U.S.), as the disease is too rare to interest for-profit drug developers, and the prevalence in the U.S. is less than 20 (8) (unpublished data from CureDRPLA’s Global Patient Registry, ClinicalTrials.gov ID NCT05489393).

Most poly-glutamine expansion disorders impact a preferential region, notably including the striatum in HD (61), and the cerebellum and brain stem in SCA1, 2, 3, 6, 7, and 17 (62).

Similarly, the spinal cord and brainstem are impacted in spinal-muscular bulbar atrophy (SBMA) (63). Unfortunately, DRPLA neuropathology is much more diffuse than these other disorders, and includes reported atrophy in the basal ganglia, brain stem, cerebellum, and cortex (64). Because of this, focally delivered treatments, such as the surgically delivered Adeno-Associated Virus gene therapies that are currently being tested in the striatum of HD patients (65), have a much higher bar to clear for DRPLA than for other diseases. Excitingly, however, recent evidence demonstrates that ASO distribution after delivery to the CSF is widespread in mice, rats, and non-human primates (66), suggesting that for the time being, ASOs remain the best option for DRPLA therapeutics focused on ATN1-lowering.

Many mouse ASO studies have been reported in other forms of CAG expansion disorders, including SBMA (67), SCA1 (68), 2 (69), 3 (70), and 7 (71). However, ours is the first to describe an ASO rescue experiment in a fully humanized mouse model, and indeed is the first report of a completely humanized mouse model of a RED, that we are aware of. We believe that complete humanization, enabled by improvements in genome editing technology (72), provides significant advantages for modeling neurodegenerative diseases. First, transgenic mouse models remain a workhorse of preclinical studies in neurodegeneration, but our approach ensures normal expression levels of the integrated gene by replacing a mouse ortholog with mutant human *ATN1*. Thus, total ATN1 protein levels, as well as the balance of WT to mATN1, are normal. Our approach enables us to screen human oligonucleotide therapies along the expanse of the human *ATN1* transcript, without confounds imposed by altered total protein levels, which is common in transgenic mouse models that overexpress target genes.

Lastly, since our model includes all intronic sequences, we can investigate phenomena such as aberrant splicing, which isn’t available in knock-in models. For example, knock-in models of mHTT, in which only exon-1 is humanized, while everything downstream is mouse sequence (73). In short, we believe that where feasible, preclinical studies of neurodegeneration should move to a humanized context.

We are struck by the severity of the phenotypes evoked by expression of *ATN1* with 112 CAG repeats. Besides the host genes of their expanded CAG repeats, our mice are isogenic with the *Htt^Q111/+^* knock-in mouse model of HD on the C57Bl/6J strain (74), which we have worked with extensively (43, 75, 76). *Htt^Q111/+^*mice are visually indistinguishable from controls throughout their lifespan and only have modest behavioral changes that do not include pronounced motor behavioral deficits (77). Similarly, knock-in SCA2 mice with 100 repeats in *ataxin-2* (*Atxn2)* have very modest phenotypes (78, 79), whereas SCA1 knock-in mice with 154 CAG repeats in *Atxn1* are profoundly impacted (all on C57Bl/6J background strain) (28). The reason for the wide difference in tolerability of these large CAG expansions in the knock-in mouse context is a compelling feature that may help explain why these diseases impact distinct clusters of cells. We speculate that this could be influenced by their subcellular localization, as ATXN1 and ATN1 are preferentially found in the nucleus, whereas HTT and ATXN2 are preferentially found in the cytoplasm (80, 81). This working model is further supported by SBMA, which is caused by CAG expansions in the androgen receptor (*AR*) gene, and primarily impacts males, in whom testosterone drives nuclear localization of AR and consequent cellular dysfunction (63). Indeed, mutations that enhance the nuclear localization of mHTT dramatically enhance its toxicity compared to the parental line in transgenic mouse models (82). This is consistent with data in both SCA1 and SCA3, in which mouse experiments reveal that preventing nuclear localization of mATXN1 and mATXN3 delays pathogenesis (83–85). We believe that our results contribute to in vivo evidence for the model that nuclear localization of expanded polyglutamine proteins is a critical driver of toxicity.

## Materials and Methods

### Mouse Development

#### Transfection of ES cells to introduce a targeted mutation via CRISPR/Cas9 mediated gene editing

The embryonic stem (ES) cells were grown on a mitotically inactivated feeder layer comprised of mouse embryonic fibroblasts in ES cell culture medium containing Leukemia inhibitory factor and FBS. The cells were co-transfected with the targeting vector and a plasmid expressing Cas9 as well as the specific sgRNA(s), the latter plasmid contains a selection cassette. One day post transfection the antibiotic was transiently added to the medium to select for transfected cells. ES cell clones were isolated as soon they show a distinct morphology and were analyzed by PCR or Southern Blotting in a primary screen.

Homologous recombinant ES cell clones were expanded and frozen in liquid nitrogen after extensive molecular validation.

#### Diploid injection

After administration of hormones, superovulated BALB/c females were mated with BALB/c males. Blastocysts were isolated from the uterus at dpc 3.5. For microinjection, blastocysts were placed in a drop of DMEM with 15% FCS under mineral oil. A flat tip, piezo actuated microinjection-pipette with an internal diameter of 12–15 µm was used to inject 10–15 targeted C57BL/6NTac ES cells into each blastocyst.

After recovery, 8 injected blastocysts were transferred to each uterine horn at 2.5 dpc, pseudopregnant NMRI females. Chimerism was measured in chimeras (G0)by coat color contribution of ES cells to the BALB/c host (black/white).

#### Revitalization of cryopreserved sperm

IVF was performed using oocytes from superovulated C57BL/6NTac females and thawed sperm from previously cryopreserved chimera spermatozoa. After overnight incubation, two cell embryos were transferred into oviducts of 0.5 dpc pseudopregnant Swiss Webster recipient females. Recipient production colony as well as post embryo transfer animals are regularly monitored for microbial contamination. Germline transmission was identified by the presence of black offspring (strain C57BL/6Ntac) and by genotyping of the black offspring via PCR to prove germline transmission of the inserted mutation.

#### Genotyping analysis

Genomic DNA was extracted from biopsies and analyzed by PCR. The following templates were used as controls: water (ctrl1), wildtype genomic DNA (ctrl2), and positive DNA sample (ctrl3).

The amplification of the internal control fragment—585 bp (ctrl)—with oligos 1260_1 and 1260_2 confirms the presence of DNA in the PCR reactions (amplification of the CD79b WT allele, nt 17714036-17714620 on Chromosome 11). The amplification of the internal control fragment—333 bp (ctrl)—with oligos 11767_3 and 11767_4 confirms the presence of DNA in the PCR reactions (amplification of the CD79b WT allele, nt 17712493-17712826 on Chromosome 11).

### Animal Care and Surgery

#### Animal Care

All mice were housed in cages of 2–5 mice per cage with corn cob bedding and on a 12-hour light-dark cycle with access to food and water *ad libitum*. Male and female mice were used in all experiments and treated equally throughout the study.

#### Neonatal ICV Injection

Based on (86), mice were anesthetized by using wet ice to induce hypothermia, which was confirmed via a gentle toe pinch and observed lack of breathing. The mice were then free hand injected with 2 µL of either hASO (30 µg), mASO (30 µg), Combo (15 µg of each ASO for a total of 30 µg), or saline using a Hamilton syringe and a 32-gauge needle. The location of the injection was two-fifths of the distance from the lambda suture of one eye with the needle being inserted 2.5–3.0 mm deep. All mice were injected on postnatal day 1–3.

#### Adult ICV Surgery

Mice were anesthetized with vaporized isoflurane and placed on a stereotaxic apparatus where the surgical plane was maintained throughout the procedure. After a small vertical incision was made on the top of the head, the mice were injected with 10 µL of either hASO (200 µg), mASO (200 µg), Combo (100 µg of each ASO for a total of 200 µg), or saline using a Hamilton syringe with a 26-gauge needle. The needle was lined up with bregma then moved laterally to the right 1.0 mm and anterior 0.3 mm with the needle being inserted 3.5–3.0 mm deep. The incision was closed with suture glue and mice were injected with 5 mg/kg of Carprofen. All surgeries were performed at five weeks of age.

### ASO Screening

#### A-431 cell culture condition

A-431 is a human epidermoid carcinoma cell line purchased from ATCC®. A-431 cells were cultured in DMEM growth medium (DMEM 10% FBS, 50 units/mL penicillin and 50 µg/mL streptomycin) at 37°C and 10% CO_2_. Cells were sub cultured or trypsinized for plating when they reached 80% confluency. Cells were trypsinized, counted and diluted to 110,000 cells per mL in room temperature growth medium before adding 100 μL of the cell suspension to the wells of collagen I coated 96-well culture plate. Immediately after plating the cells, 11 μL of 10X ASO in water was added to the appropriate wells. The culture plate was incubated in a humidified incubator at 37°C and 10% CO_2_. After 48 hours, the cells were washed once with PBS before lysing with guanidine isothiocyanate-containing buffer for RNA isolation and analysis. For each treatment condition duplicate wells were tested.

#### mRNA extraction and qPCR

Each cortex and spinal cord sample was homogenized in 1 mL of Trizol reagent (Thermofisher scientific, Waltham, MA) and total RNA was extracted using the Life Technologies mini-RNA purification kit (Qiagen, Valencia, CA) according to the manufacturer’s protocol. After purification, the RNA samples were subjected to real-time RT-PCR analysis using the Life Technologies ABI QuantStudio 7 Flex Sequence Detection System (Applied Biosystems Inc, Carlsbad, CA). Briefly, 10 µL RT-PCR reactions containing 400 nL of RNA were run with the AgPath-ID One-Step qRT-PCR Kit (Thermofisher scientific, Waltham, MA) reagents and the primer probe sets. This system uses real-time florescence PCR to quantitatively determine mRNA expression levels. The assay is based on a target-specific probe labeled with a fluorescent reporter and quencher dyes at opposite ends. The probe is hydrolyzed through the 5‘-exonuclease activity of Taq DNA polymerase, leading to an increasing fluorescence emission of the reporter dye that can be detected during the reaction. All qPCR reactions were run in triplicate. Outliers in triplicates due to technical issues were eliminated. Target mRNA was then normalized to *Ppia*, a ubiquitously expressed housekeeping gene, and this was further normalized to the level measured in control animals that were administered PBS. mRNA levels are reported as percent control.

### Behavior

#### Balance Beam

The balance beam apparatus consisted of a metal rod (length: 100 cm, diameter: 12 mm and 6 mm) set up at a tapered angle with a dark box at one end and 60-watt lamp on the other. The dark box had nestling and bedding from the mouse’s home cage. The dark box and rod were cleaned with 70% ethanol between cages. Mice were trained for two days with three trials on both of the rods. The mice were tested on the third day with three trials on the 6 mm rod. All mice were tested and trained from 6–7 weeks of age.

#### Fixed Rotarod

Motor coordination was tested using a fixed rotarod apparatus (Harvard Apparatus, 76-0770). Each mouse went through three 5-minute training trials at two speeds (4-and 8-rpm). Mice were then tested for three trials at both speeds with a max time of 5 minutes. All mice were tested and trained from 6–7 weeks of age.

#### Open Field

Mice were placed in the center of a 41 x 41 cm closed box with a transparent acrylic lid. A GoPro camera, mounted in the lid, recorded a 12-minute video of each individual mouse. Quantitative analysis was performed using Cai Lab’s EzTrack script to measure total distance travelled and the freezing to motion ratio (87). Open field data presented here is from a cohort of mice from a later study that utilized the same mouse model and hASO treatment, however the mice did receive multiple intraperitoneal injections of saline (Table S3).

#### Modified SHIRPA

This screening test is a modified version of (88) SHIRPA assessment (**S**mithKline Beecham Pharmaceuticals; **H**arwell, MRC Mouse Genome Centre and Mammalian Genetics Unit; **I**mperial College School of Medicine at St Mary’s; **R**oyal London Hospital, St Bartholomew’s and the Royal London School of Medicine; **P**henotype **A**ssessment). In our modified SHIRPA we observed mice inside of their home cage to assess their level of lethargy, gait issues, tremors, pelvic and tail elevation, hindlimb and forelimb grasping, piloerection, palpebral closure, and touch escape. Each phenotype that was assessed had an individual scale that was used to measure the behavior (Table S5). Overall, the higher the mouse’s score for any phenotype equated to worsening of the phenotype in the mouse. All mice were assessed at 4 weeks of age.

#### Analysis of Circadian Patterns of Home Cage Behavior

A subset of male *Atn1^Q112/+^* mice (n = 4) aged eight weeks old were followed in a behavioral activity monitored home cage using the Noldus PhenoTyper system (Noldus Information Technology, Wageningen, The Netherlands) as described previously (89). Plexiglas cages measuring 30 x 30 cm included corncob bedding, with cameras recording overhead for the entirety of the monitoring period. All cages were recorded simultaneously for 67-hours, beginning at 12 p.m. on Day 1 and ending at 7 a.m. Mice were able to freely enter a plastic enclosure (termed ‘hidden shelter’) with two entry/exit points.

EthoVision XT software (Noldus Information Technology, Wageningen, The Netherlands) was used to track the movement of the mice for the duration of the experiment. Utilizing this software, the 67-hours of behavioral tracking data was analyzed offline in an automated manner by a blinded investigator by dividing the cage recording into various zones (arena center, arena perimeter, hidden shelter) and the duration within each zone (seconds) determined for each mouse over the monitoring period. Relative measurements of the arena areas were as follows: arena center 293 cm^2^, arena perimeter 347 cm^2^, and hidden shelter 106 cm^2^. Behavioral activity measurements were binned into 1-hour sessions and activity tracked across all zones, with average activity of each bin taken for the light periods (Light-1,-2, and-3) in comparison with the dark periods (Dark-1,-2, and-3). Two saline-treated *Atn1^Q112/+^*mice died during the monitoring period (at 31-and 47-hour time points) and behavioral analysis until death was included for analysis; all other animals survived the entirety of the monitoring period. Mice were 8 weeks of age during testing. Total activity for each 1-hour bin and average activity in each light phase were analyzed by mixed-effects model (time x treatment), with p < 0.05 considered significant (using GraphPad Prism version 9.0 or later).

### ATN1 and NEFL quantification

#### RNA Extraction and cDNA Synthesis

Cortical tissue (30–50 mg) was homogenized in 500 µL of QIAzol Lysis Reagent using the Bead Blaster 24 (Benchmark). RNA was extracted using the RNeasy Lipid Tissue Mini kit (Qiagen 74804) according to the manufacturer’s instructions. RNA quality was tested using the NanoDrop Spectrophotometer 2000 (Thermo Fisher Scientific, ND-2000). cDNA synthesis was carried out using the SuperScript III First-Strand Synthesis System (Thermo Fisher Scientific) according to the manufacturer’s instructions.

#### RT-qPCR

RT-qPCR used 10 µL TaqMan Universal Master Mix II with UNG (Thermo Fisher Scientific, 4440046) along with 5 µL molecular biology grade water, 3 µL cDNA, and 2 µL of probes. The reaction was quantified on the QuantStudio 7 Flex Real-Time PCR System (Thermo Fisher Scientific, 4485688). β-Actin (Thermo Fisher, Mm02619580_g1) was used as the reference gene to determine the relative expression of ATN1 (Thermo Fisher Scientific, Hs01073465_m1), and Atn1 (Thermo Fisher Scientific, Mm01185029_m1). This expression was compared to the average expression of control samples. Fold change was given by 2^(-ΔΔCt)^.

#### NEFL Quantification

NEFL was quantified using an electrochemiluminescent ELISA measured with a MESO QuickPlex SQ120MM (Meso Scale Discovery; MSD). 96 well plates (MSD, L45SA-1) were coated in capture antibody (MSD, F217X-3) that was diluted in Diluent 100 (MSD, R50AA-2) and incubated for one hour at room temperature with shaking at 700-rpm. The plates were rinsed with wash buffer (PBS with 0.2% tween-20) three times. NEFL calibrator (MSD, F217X-3) was serially diluted in Diluent 12 (MSD, R50JA-2) and plasma samples were diluted at a 1:10 dilution in Diluent 12. The diluted calibrator and samples were then loaded into the plate(s) and incubated at room temperature for one hour with shaking at 700-rpm. Plates were rinsed again with wash buffer three times then coated in sulfo-tag conjugated detection antibody (MSD, F217X-3) that was diluted to 1x with Diluent 11 (MSD, R55BA-3). Plates were incubated at room temperature for one hour while shaking at 700rpm then rinsed again with wash buffer three times.

Finally, 150 µL of Read Buffer B (MSD, R60AM-1) was added to the plates which were then read with MESO QuickPlex SQ120MM.

### Neuropathology

#### Tissue Preparation and histochemical staining

At nine weeks of age mice were sacrificed via CO_2_. Half of the brain was drop fixed in 10% non-buffered formalin. Hemi-brains were removed after 12 hours and transferred into 1x PBS with 0.02% sodium azide.

Parasagittal sections were prepared from 32 fixed hemibrains by embedding in paraffin and sectioning at 5 µm along the sagittal plane using a standard microtome to the level of the mid-olfactory bulb. Four brains were mounted on each slide for further evaluation. H&E staining was performed on several of these sections according to standard protocol.

#### Immunohistochemistry

Immunohistochemistry was performed on parasagittal brain sections using a Leica Bond III Fully Automated IHC and ISH Staining System (Leica Biosystems, Wetzlar, Germany). The sections were immunostained with mouse monoclonal antibody against glial fibrillary acidic protein (GFAP, 1:600; Biocare Medical CM065A), Iba1 (1:1000; Wako 019-19741), NeuN (1:500; EMD Millipore MAB377), and p62 (1:500; Abcam ab56416). Appropriate positive and negative controls were included with each antibody and each run.

### Bulk RNA Sequencing

cDNA libraries were constructed at Azenta (South Plainfield, NJ) using the Illumina TruSeq RNA Sample Prep Kit with ERCC spike-in and sequenced on a HiSeq 2000 (2 x 150 bp) to an average read depth of 2.9e7. Sequence reads were trimmed to remove adapter sequences and nucleotides with poor quality using Trimmomatic v.0.36. Reads were aligned to the *Mus musculus* GRCm38 ERCC reference genome assembly (ENSEMBL) using STAR aligner v.2.5.2b. Unique gene hit counts were calculated using `featureCounts` from the Subread package v.1.5.2. Differential gene expression analysis was conducted in R using `edgeR` (90), with `voom` (91). Unsupervised clustering was performed using package `pheatmaps`. For correlation plots in Fig. 6B, Log2FC for the indicated comparison was converted to z-scores for each gene. The GO enrichment plots from Fig. S13A were generated with package `clusterProfiler` (92).

### Statistics

Behavioral and molecular data were recorded and stored in a Carroll lab cloud storage solution (Google Drive). Raw data was read into an R project for processing, statistical comparisons and generating graphs, which were finalized in Adobe Illustrator. Analysis of molecular and behavioral data was done using one or multi-factor ANOVAs, with Tukey’s Honest Significance posthoc tests. Count data (e.g. SHIRPA) was analyzed in R using a general linear model (GLM) with a Poisson log-link function. All R scripts used for analysis and figure generation are available on the linked Dryad data repository.

### Study Approval

All animal experimentation was approved by the University of Washington’s Institutional Animal Care and Use Committee.

## Data Availability

RNAseq results are available at gene expression ombibus (GEO). All other raw data and R scripts for analysis are available at a Dryad repository (https://datadryad.org), searchable by the name of this manuscript.

---

**Supplemental Figure 1:**
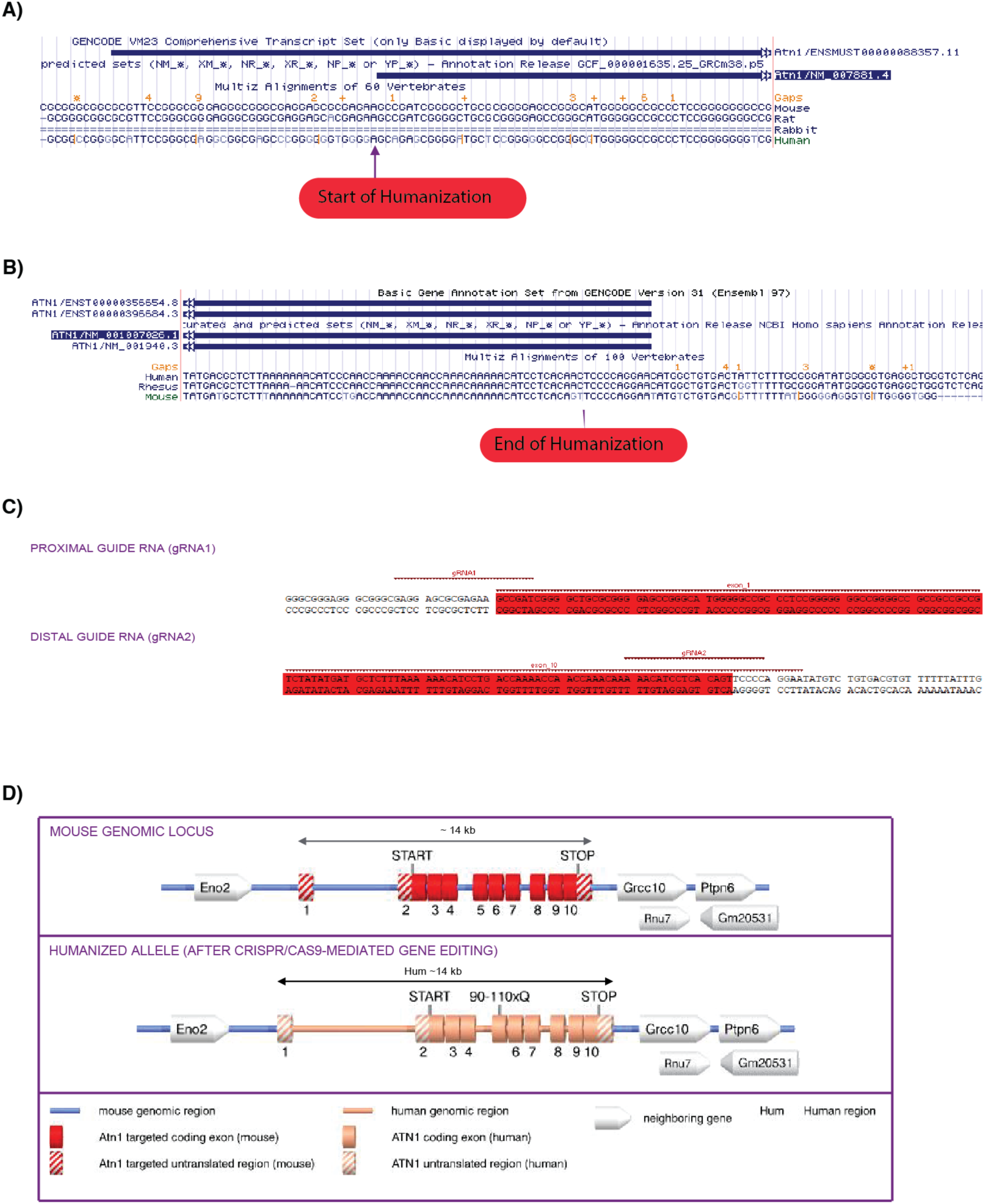
Summary of approach to generating *Atn1^Q112/+^*mouse generation a fully humanized *ATN1* allele. Detailed view of the genomic location of the beginning (A) and end (B) of humanization in the mouse genome. C) gRNA sequences used for engineering the *Atn1* allele in vitro. D) Cartoon summary of the humanized *ATN1* allele including neighboring genes (not to scale).

**Supplemental Figure 2:**
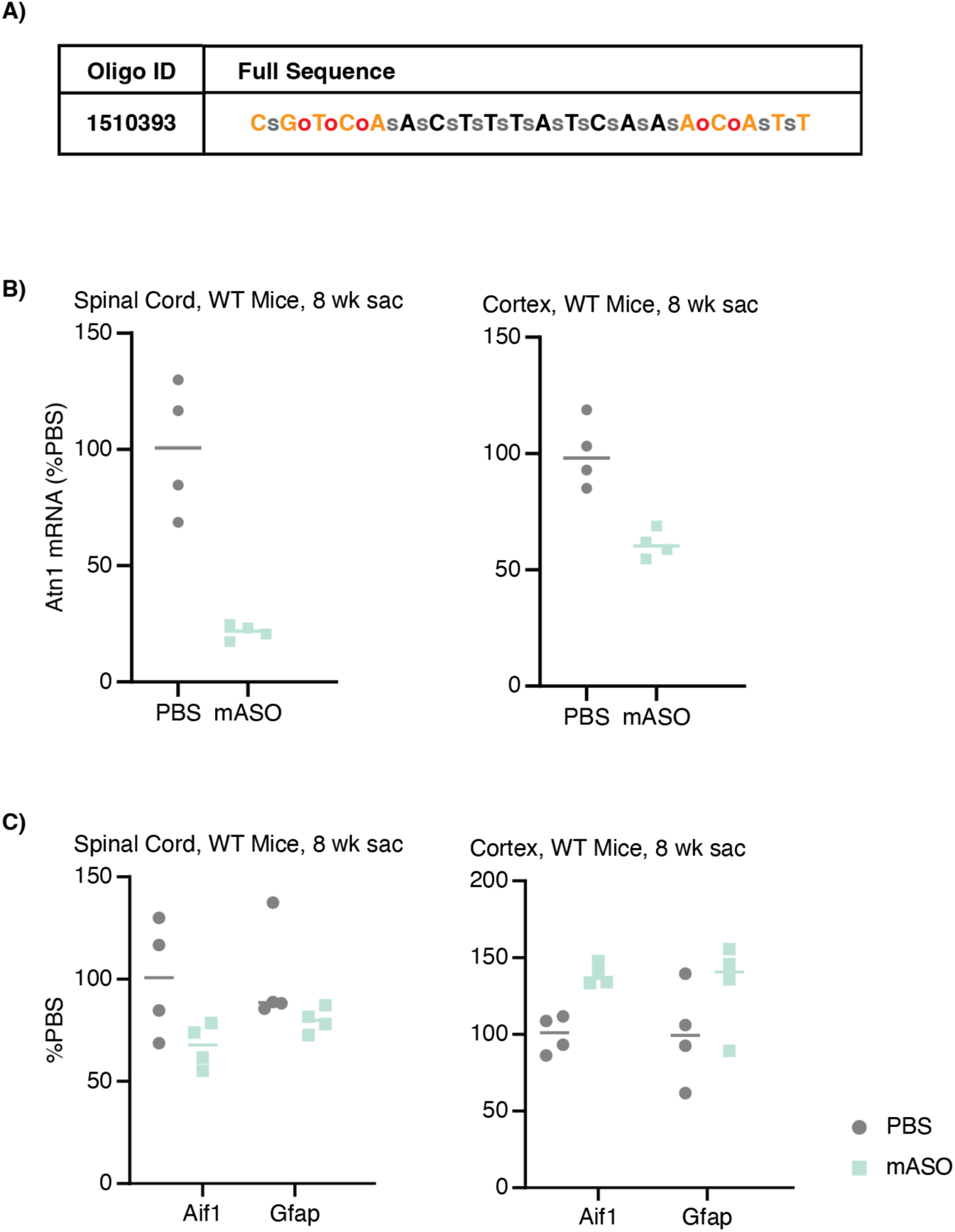
Ionis *in vivo* mouse ASO sequence with toxicity and tolerability tests. **A)** mASO (Ionis ID # 1510393) sequence. Orange indicates a nucleotide with 2’MOE modification, all the “C” are 5MeC. Black is DNA. Red “o” is phosphate backbone, and “s” is phosphothioate backbone. **B)** WT C57BL/6NT mice dosed with 700 µg mASO led to sufficient knockdown of Atn1 in cortex and spinal cord 8-weeks post dosing. No deaths were seen acutely or longitudinally indicating a lack of toxicity. **C)** Glial activation was quantified via qPCR by looking at Aif1 and Gfap in cortex and spinal cord in WT mice 8-weeks post 700 µg injection. No abnormalities were detected.

**Supplemental Figure 3:**
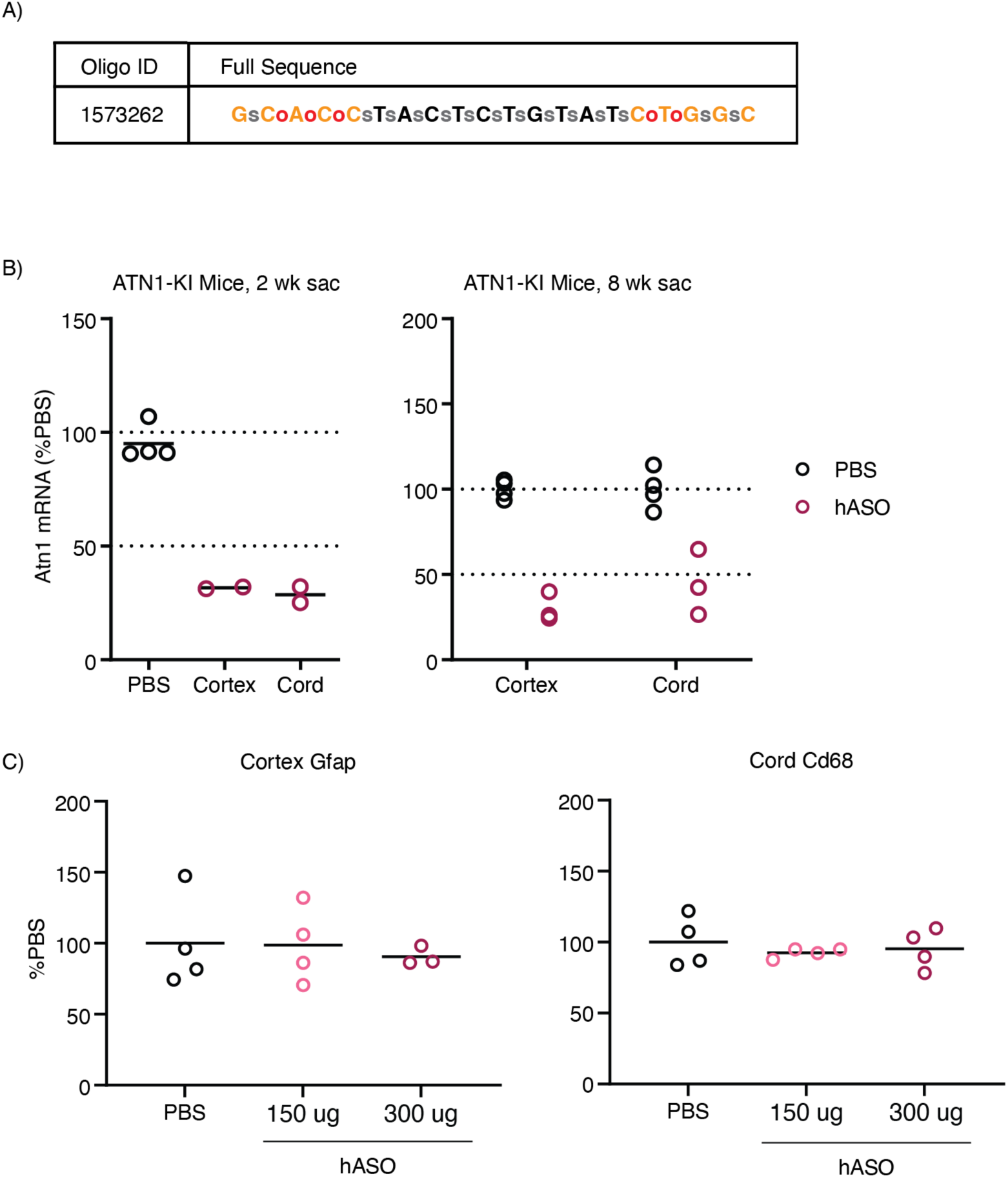
Ionis *in vivo* human ASO sequence with toxicity and tolerability tests. **A)** hASO (Ionis ID # 1573262) sequence. Orange indicates a nucleotide with 2’MOE modification, all the “C” are 5MeC. Black is DNA. Red “o” is phosphate backbone, and “s” is phosphothioate backbone. **B)** Knockin C57BL/6NT ac-Atn1em7219 mice dosed with 300 µg hASO led to sufficient knockdown of ATN1 in cortex and spinal cord at 2-and 8-weeks post dosing. No deaths were seen acutely or longitudinally indicating a lack of toxicity. **C)** WT mice were treated with hASO at 150 µg and 300 µg. Glial activation was quantified via qPCR by looking at Gfap in cortex and Cd68 in spinal cord in WT mice 2-and 8-weeks post injection. No abnormalities were detected.

**Supplemental Figure 4:**
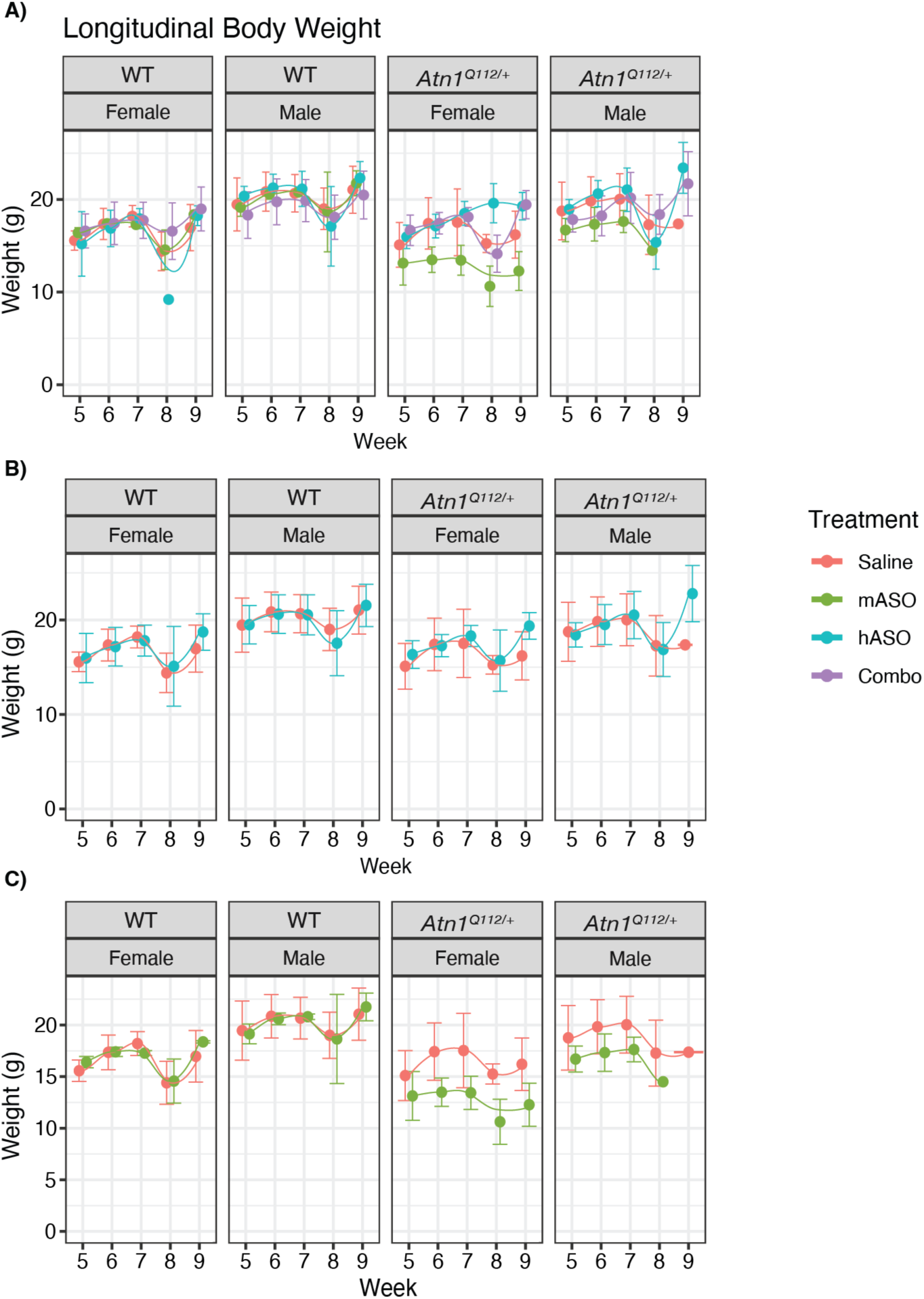
Longitudinal body weight A-C) Line plots showing longitudinal body weight, measured in grams, from 5, 6, 7, 8, and 9 weeks of age. N = 102, 100, 88, 56, and 59 for each respective week. Each plot is facetted by genotype and sex with A showing all the treatment arms, B showing only saline and hASO arms, and C showing only saline and mASO arms.

**Supplemental Figure 5:**
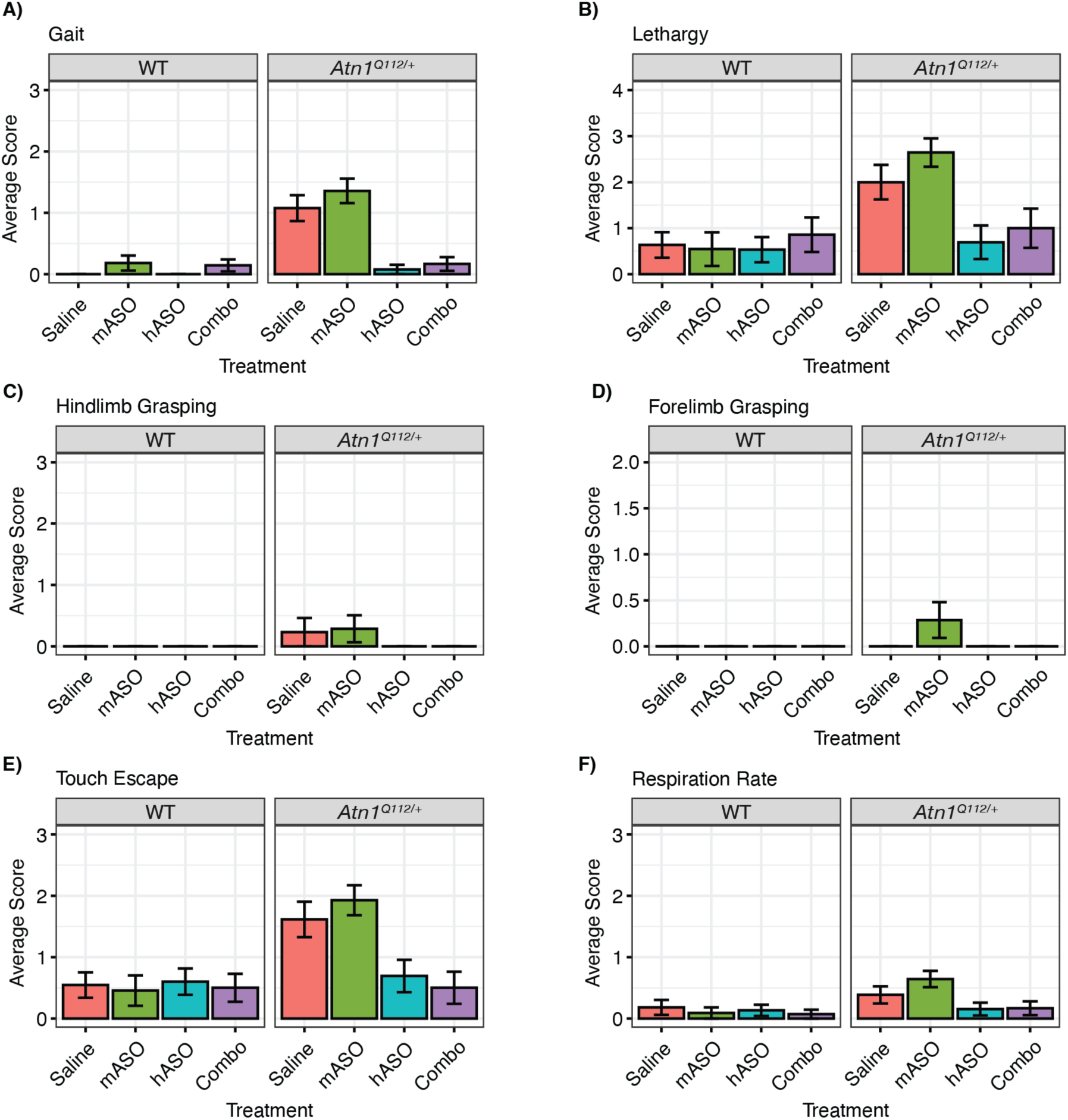
Human, but not mouse, *ATN1* ASO treatment improves SHIRPA scores in *Atn1^Q112/+^* mice. A-E) Representative bar plots displaying the mean of the observed phenotype with error bars indicating the standard error. HASO-treated *Atn1^Q112/+^* mice showed an overall reduction in gait (A), lethargy (B), touch escape (E), and respiration rate (F). There was no significant observation of hindlimb (C) or forelimb grasping (D) across all arms.

**Supplemental Figure 6:**
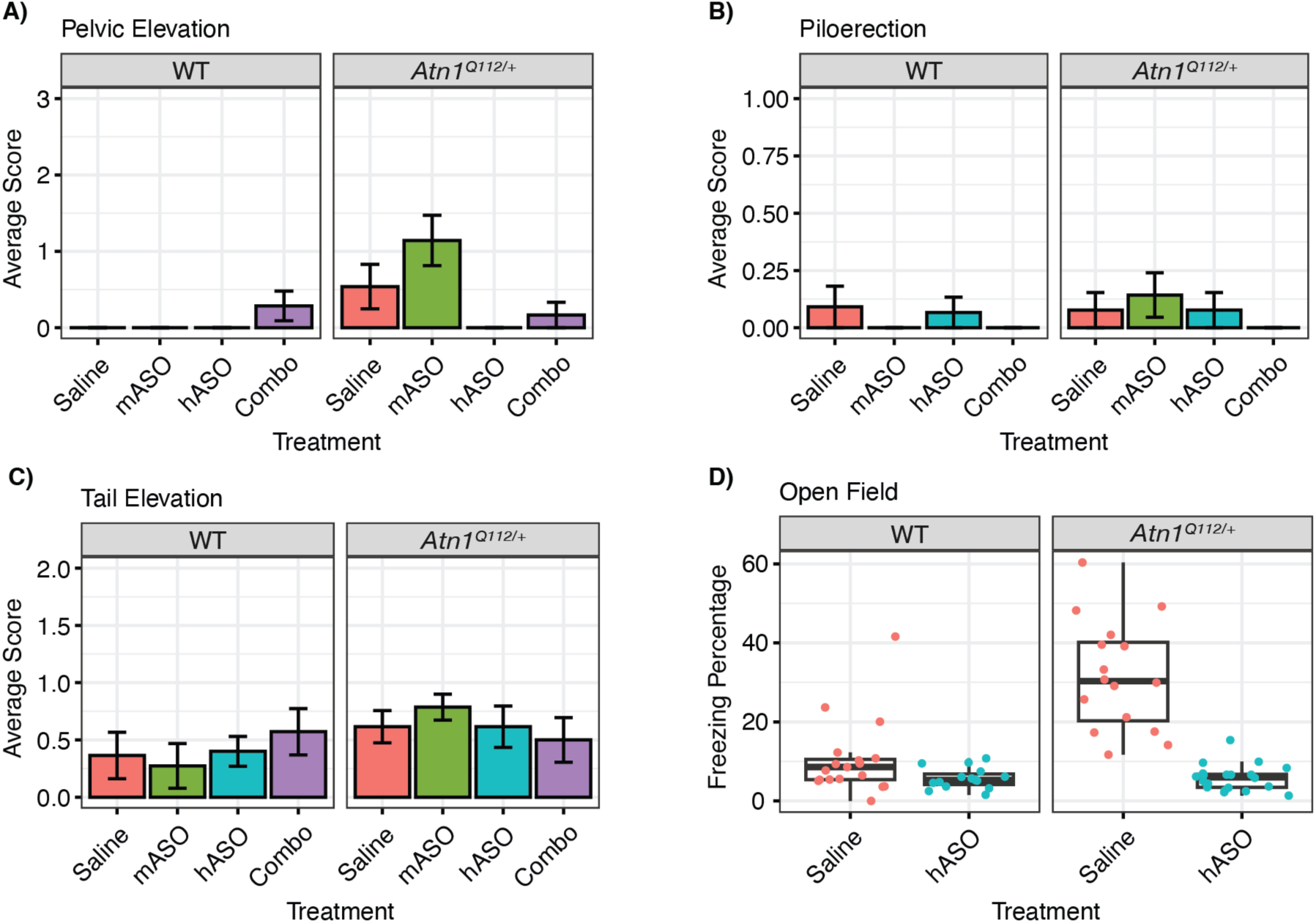
Human, but not mouse, *ATN1* ASO treatment improves SHIRPA scores in *Atn1^Q112/+^* mice. A-C) Representative bar plots displaying the mean of the observed phenotype with error bars indicating the standard error. *Atn1^Q112/+^* mice treated with hASO showed an overall reduction in pelvic elevation (A) and piloerection (B). However, *Atn1^Q112/+^*mice treated with hASO did not see any improvement in tail elevation issues (C). **D)** Boxplot showing freezing percentage during an open field test in hASO-and saline-treated WT and *Atn1^Q112/+^* mice. *Atn1^Q112/+^*mice treated with hASO have an overall lower freezing percentage than controls.

**Supplemental Figure 7:**
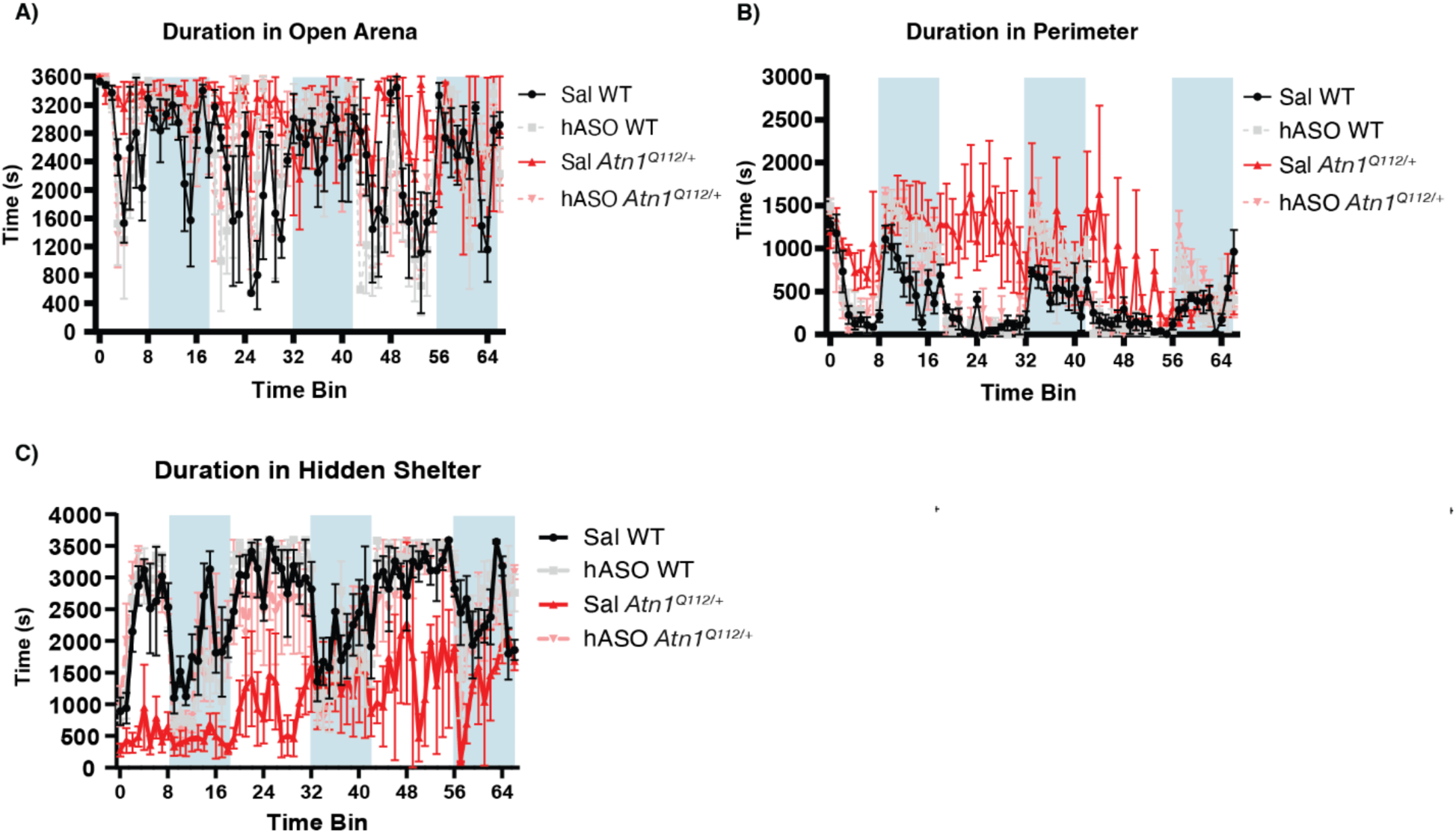
Treatment with human ASO rescues circadian behavioral deficits in *Atn1^Q112/+^* mice. All analyses measured by mixed-effects model and Tukey’s post-hoc test (n = 4/group, with n = 2 *Atn1^Q112/+^* mice dying in the last sessions). **A)** Duration in the entire chamber during each 1-hour bin for a 67-hour recording session. There was a significant time x treatment group interaction (F_(198, 742)_ = 1.277, p = 0.013), with post-hoc tests demonstrating marked differences in arena exploration time in saline-treated *Atn1^Q112/+^* mice versus saline-treated WT mice at the 26-and 30-hour time bins (p < 0.05; *not shown*). **B)** Duration in perimeter zone during each 1-hour bin for a 67-hour recording session. There was a significant time x treatment group interaction (F_(198, 737)_ = 1.797, p < 0.0001), with post-hoc tests demonstrating marked differences in total time spent in the perimeter zone for saline-treated *Atn1^Q112/+^* mice versus saline-treated WT mice at the 3-, 6-and 8-hour time bins (p < 0.05; *not indicated for clarity*). Notably, post-hoc tests also revealed marked differences in total time spent in the perimeter zone for saline-treated *Atn1^Q112/+^* mice versus hASO-treated *Atn1^Q112/+^* mice at the 2-, 3- and 57-hour time bins (p < 0.05; *not indicated for clarity*). **C)** Duration in hidden shelter during each 1-hour bin for a 67- hour recording session. There was a significant time x treatment group interaction (F_(198, 737)_ = 1.486, p < 0.0001), with post-hoc tests demonstrating marked differences in total time spent in the hidden shelter for saline-treated *Atn1^Q112/+^* mice versus saline-treated WT mice at the 2-, 3-, 7-, 8-, 10-, 15-, 18-, 19-, 20-, 24-, 27-, 29-, 30-, 43-, and 56-hour time bins (p < 0.05; *not indicated for clarity*). There were also marked post-hoc differences in total time spent in the hidden shelter for saline-treated *Atn1^Q112/+^* mice versus hASO-treated *Atn1^Q112/+^* mice at the 2-, 3-, 5-, 6-, 7-, 8-, and 57-hour bins (p < 0.05; *not indicated for clarity*).

**Supplemental Figure 8:**
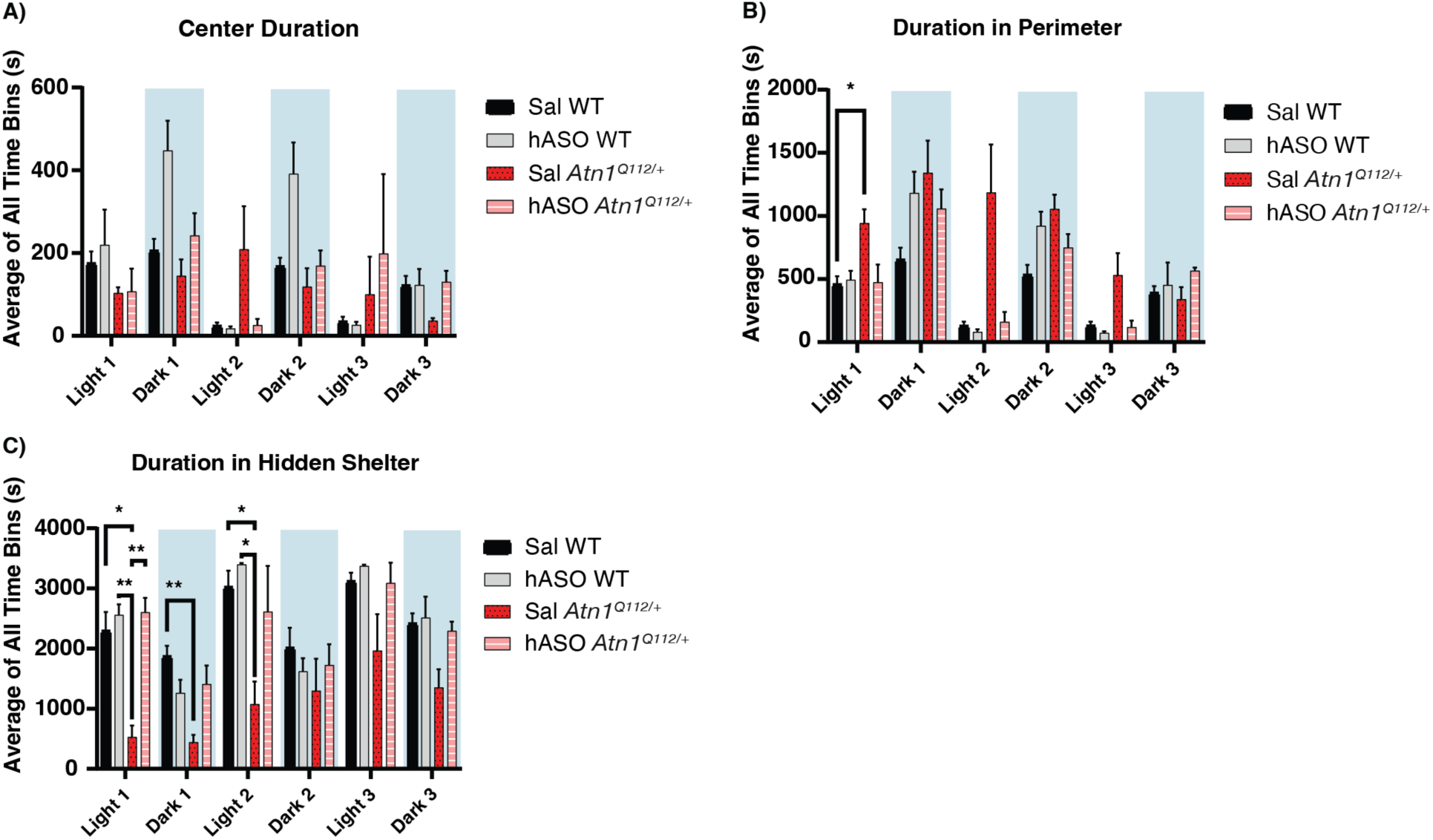
Human *ATN1* ASO treatment rescues circadian behavioral deficits in *Atn1^Q112/+^* mice. The average activity of all time bins during the light and dark phases of home cage monitoring also reveals marked impacts of hASO administration in *Atn1^Q112/+^* mice. All analyses measured by mixed-effects model and Tukey’s post-hoc test (n = 4/group, with n = 2 *Atn1^Q112/+^*mice dying in the last sessions) **A)** Average time elapsed in the chamber center during each 1-hour bin for the light versus dark periods during a 67-hour recording session demonstrates a significant time x treatment interaction (F_(15, 55)_ = 2.973, p < 0.002), with no post-hoc differences between groups. **B)** Average time elapsed in the home cage chamber perimeter zone during each 1-hour bin for the light versus dark periods during a 67-hour recording session showed a significant time x treatment interaction during the monitoring period (F_(15, 56)_ = 2.473, p < 0.008). Post-hoc analysis of perimeter exploration time demonstrated that hASO treatment in *Atn1^Q112/+^* mice normalized behavior in the first light phase of recording (* p < 0.05). **C)** Average time elapsed in the chamber hidden shelter during each 1-hour bin for the light versus dark periods during a 67- hour recording session shows a significant main effect of treatment group (F_(3, 12)_ = 9.274, p < 0.002). Saline-treated *Atn1^Q112/+^*mice show significant delays in spending time in the hidden shelter, indicative of significant behavioral disruptions that can be rescued with hASO treatment, as demonstrated by post-hoc tests with *p < 0.05.

**Supplemental Figure 9:**
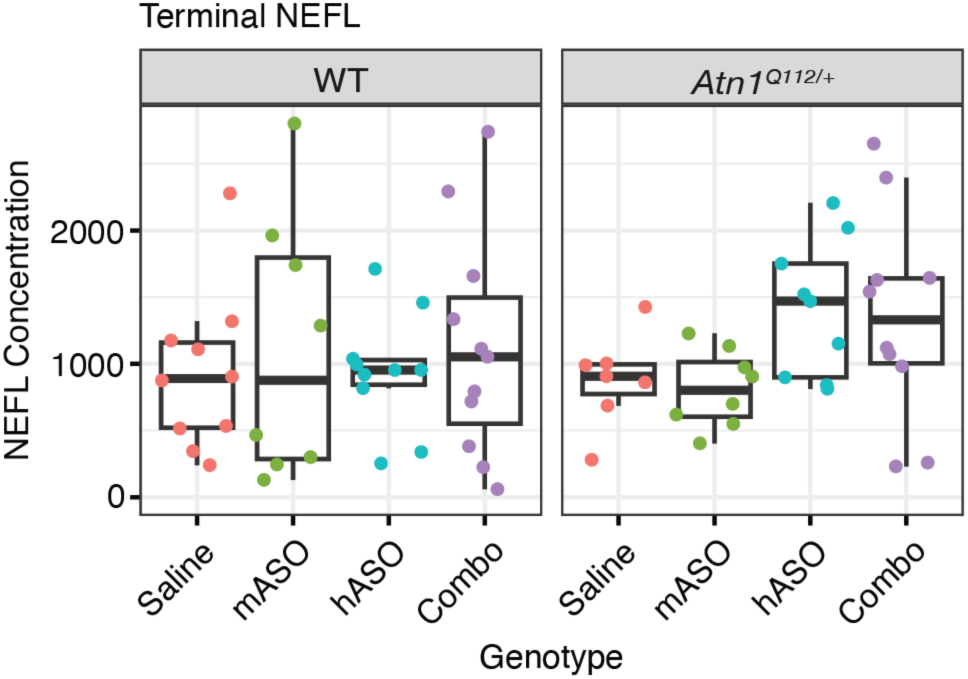
NEFL. Terminal NEFL was measured from plasma (n = 73). There was no observed significant change in any WT or *Atn1^Q112/+^* mice in any treatment arms.

**Supplemental Figure 10:**
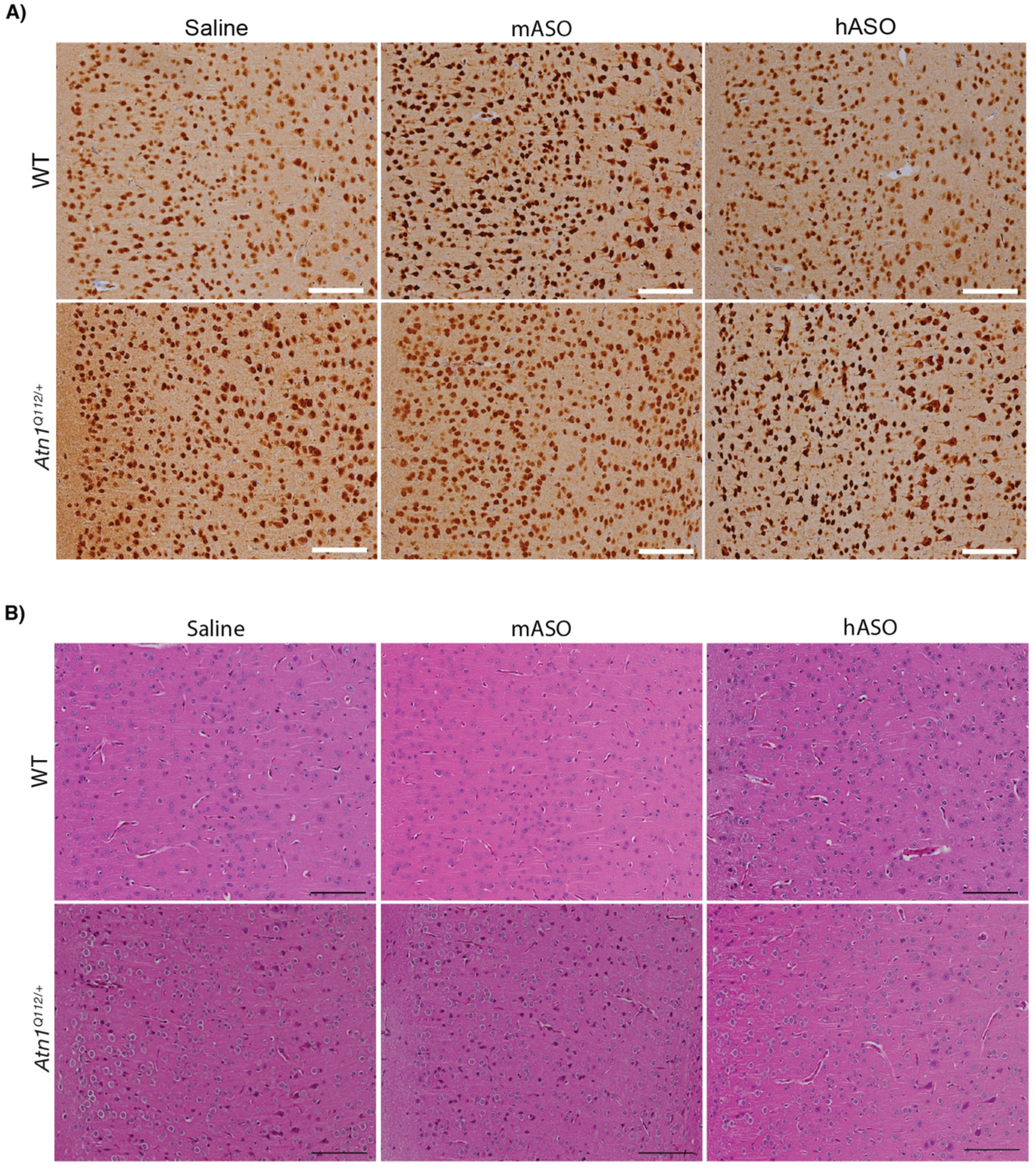
Immunohistochemical and histochemical evaluation for neurons and neuropil integrity shows no evidence of neuron loss in *Atn1^Q112/+^* mice compared to WT littermates in any treatment group. **A)** Representative light microscopy images of frontal cortex in parasagittal sections, selected from each genotype/treatment group and stained for NeuN by IHC. Scale bars = 100 µm. **B)** Representative light microscopy images of frontal cortex in parasagittal sections, selected from each genotype/treatment group and stained with hematoxylin and eosin to evaluate neuropil integrity and evidence of microvacuolization. Scale bars = 100 µm.

**Supplemental Figure 11:**
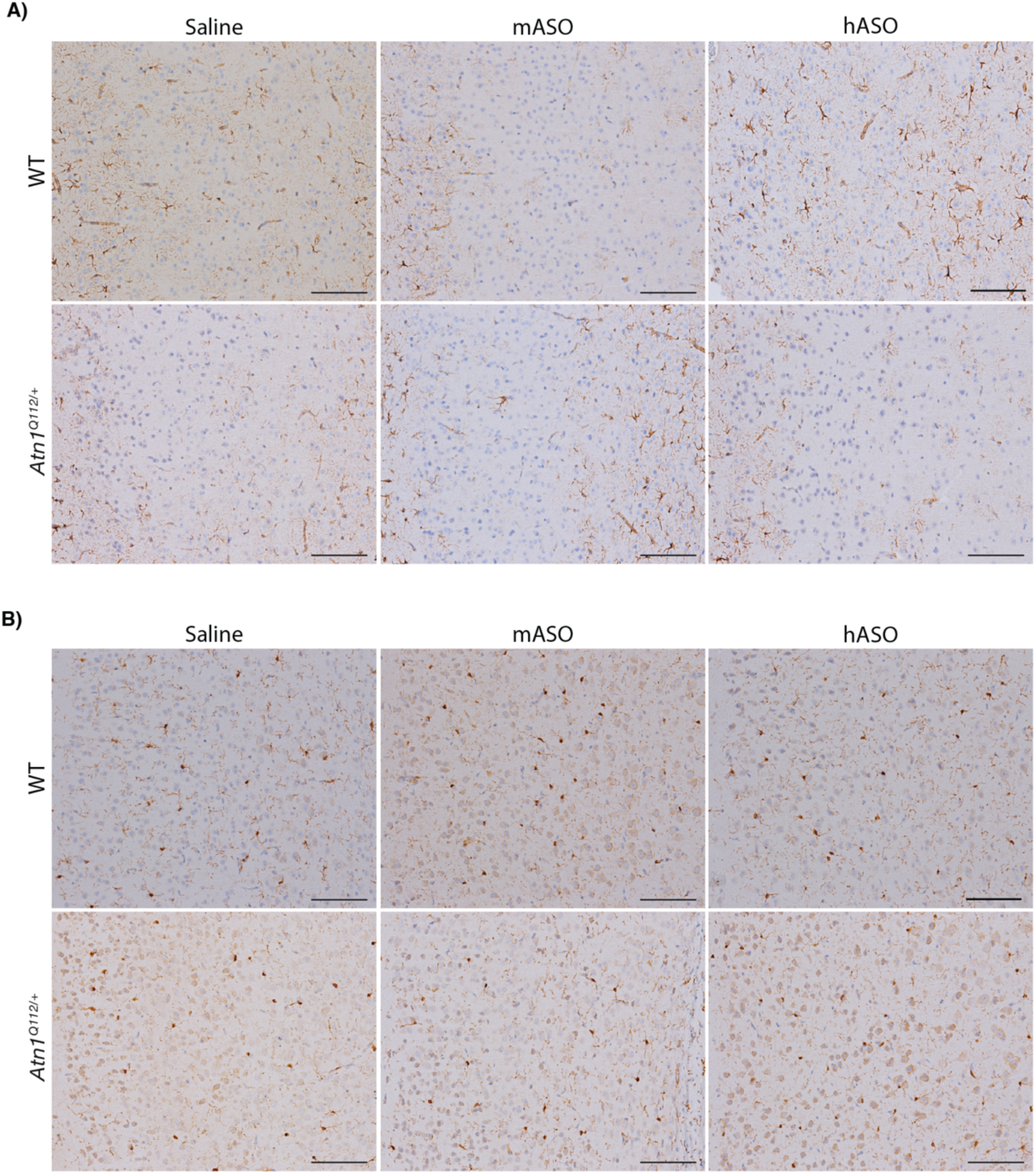
Immunohistochemical evaluation for glial cells shows no evidence of reactive gliosis in *Atn1^Q112/+^* mice compared to WT littermates in any treatment group. **A)** Representative light microscopy images of frontal cortex in parasagittal sections, selected from each genotype/treatment group and stained for GFAP by IHC to evaluate astrocytes and assess for reactive gliosis. Scale bars = 100 µm. **B)** Representative light microscopy images of frontal cortex in parasagittal sections, selected from each genotype/treatment group and stained with Iba1 to evaluate microglia and assess for microglial nodules. Scale bars = 100 µm.

**Supplemental Figure 12:**
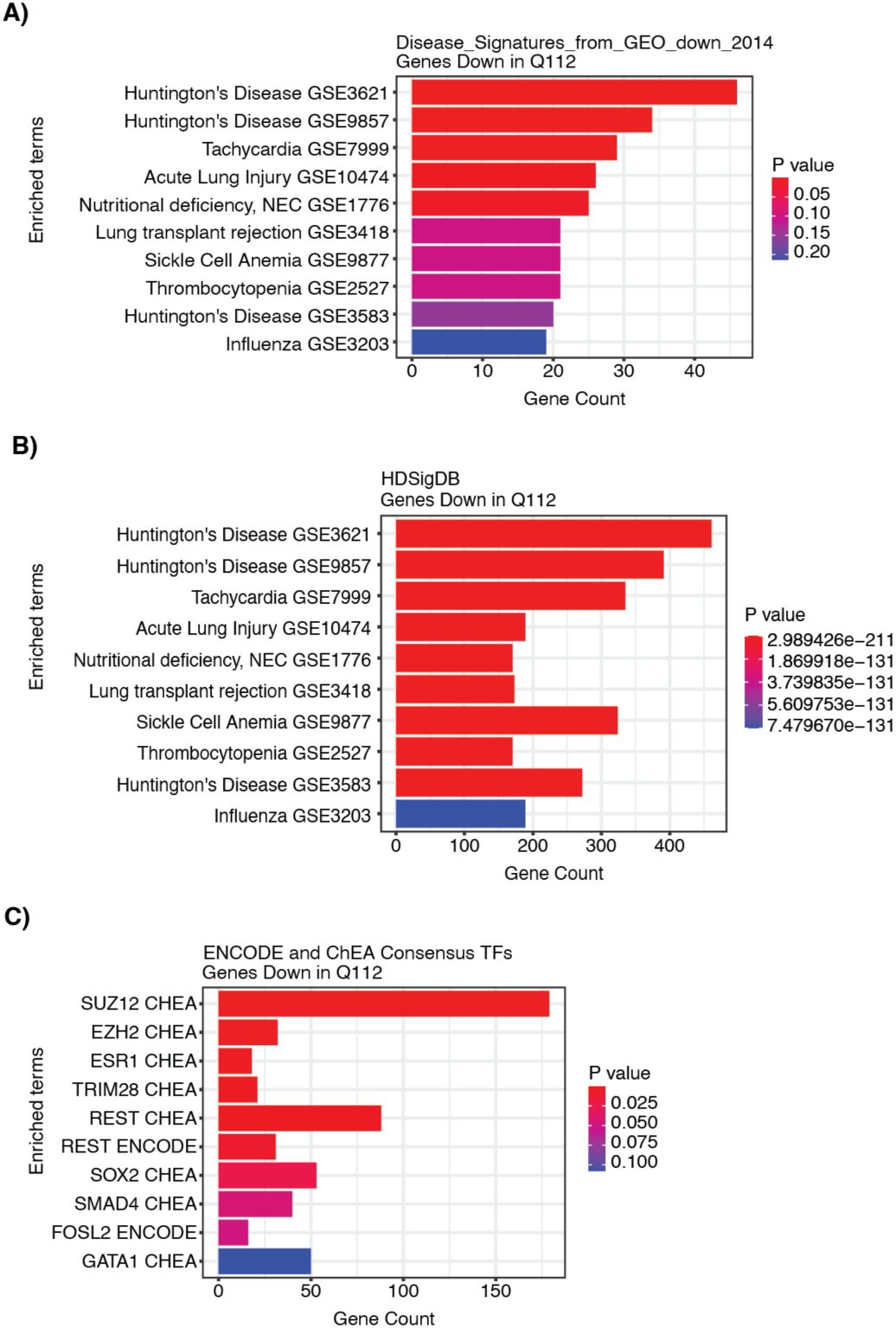
Pathway analysis on genes down-regulated in *Atn1^Q112/+^* cerebellum suggests shared transcriptionopathy with Huntington’s disease. Genes identified as down-regulated in the cerebellum of *Atn1^Q112/+^* mice versus WT at a threshold of FDR < 0.05 and Log2FC <-0.58 (1041 genes) were used to input to Enrichr. **A)** Down-regulated genes were assessed for disease signatures in GEO datasets, finding significantly enriched in two associated with transcriptional deficits in Huntington’s disease (GSE3621, 46/300 overlap, padj = 5.14e-9; and GSE9857, 34/300 overlap, padj = 0.001). **B)** Using this same set of 1041 down-regulated genes as input for the HDSigDB Enrichr library, we detected significant overlap with datasets characterizing cerebellar dysregulation (e.g. down-regulated genes in the 10 month-old Q175 mouse model of HD versus controls, GSE73468, 461/1870 overlap, padj = 7.03e-208; genes down-regulated in the cerebellum of N171-82Q N-terminal model of HD at 10 weeks of age, 391/1469 genes, padj = 9.81e-184). **C)** Known transcription factor targets were assessed using the ENCODE and ChEA consensus library, finding that the top enrichment hits were both PRC2 subunits (SUZ12 CHEA, 179/1684 overlap, padj = 3.10e-19; EZH2 ChEA, 32/237 overlap, padj = 3.55e-5).

**Supplemental Figure 13:**
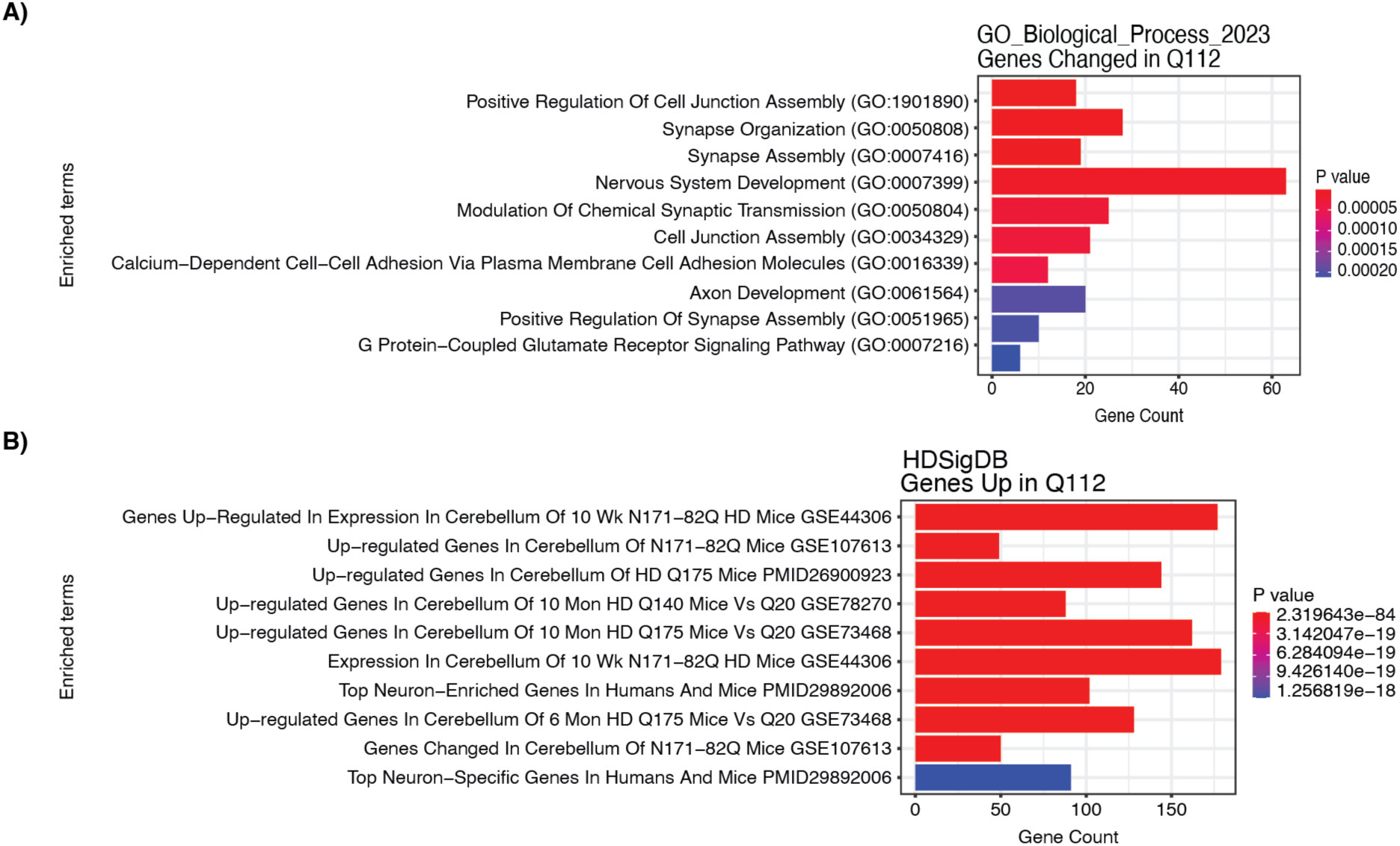
Pathway analysis on genes up-regulated in *Atn1^Q112/+^* cerebellum highlights up-regulation of neuronal genes and shared molecular pathology with Huntington’s disease. Genes identified as up-regulated in the cerebellum of *Atn1^Q112/+^* mice versus WT at a threshold of FDR < 0.05 and Log2FC > 0.58 (655 genes) were used to input to Enrichr **A)** Gene Ontology Biological Process terms for genes associated with neuronal function are enriched (e.g. Calcium-Dependent Cell-Cell Adhesion Via Plasma Membrane Cell Adhesion Molecules GO:0016339, 11/38 overlap, padj = 3.28e-5; Synapse Organization GO:0050808, 19/131 overlap, padj = 3.79e-5). **B)** *Atn1^Q112/+^* cerebellar up-regulated genes are enriched in genesets with up-regulated genes from Huntington’s disease mouse models (e.g. Genes Up-Regulated In Expression In Cerebellum Of 10 Wk N171-82Q HD Mice GSE44306, 177/976 overlap, padj = 4.95e-84; Up-regulated Genes In Cerebellum Of 10 Mon HD Q140 Mice Vs Q20 GSE78270, 88/600 overlap, padj = 1.59e-30).

**Supplemental Table 1:** IVF dams received from Taconic Biosciences. Details of the dams we received from Taconic Biosciences post IVF along with the birth date and size of the litter they produced. All mouse litters born from dams that arrived on September 9^th^, 2024, were only used in the open field assessment.

| # of Dams | Sex | Color | Week of Dam Birth | Arrival Date | Litter Birth Date | Litter Size |
| --- | --- | --- | --- | --- | --- | --- |
| 2 | F | Albino | 3/20/2023 | 5/17/2023 | 5/22/2023 | 11 |
| 1 | F | Albino | 3/20/2023 | 5/17/2023 | 5/23/2023 | 11 |
| 2 | F | Albino | 3/20/2023 | 5/17/2023 | 5/23/2023 | 5 |
| 4 | F | Albino | 3/20/2023 | 5/17/2023 | 5/23/2023 | 6 |
| 4 | F | Albino | 3/20/2023 | 5/17/2023 | 5/23/2023 | 7 |
| 1 | F | Albino | 3/20/2023 | 5/17/2023 | 5/23/2023 | 8 |
| 1 | F | Albino | 3/20/2023 | 5/17/2023 | 5/23/2023 | 9 |
| 2 | F | Albino | 3/20/2023 | 5/17/2023 | 5/23/2023 | 10 |
| 1 | F | Albino | 3/20/2023 | 5/17/2023 | 5/23/2023 | 12 |
| 1 | F | Albino | 3/20/2023 | 5/17/2023 | 5/23/2023 | 13 |
| 1 | F | Albino | 3/20/2023 | 5/17/2023 | 5/23/2023 | 14 |
| 1 | F | Albino | 7/8/2024 | 9/5/2024 | 9/9/2024 | 3 |
| 1 | F | Albino | 7/8/2024 | 9/5/2024 | 9/9/2024 | 4 |
| 5 | F | Albino | 7/8/2024 | 9/5/2024 | 9/9/2024 | 6 |
| 1 | F | Albino | 7/8/2024 | 9/5/2024 | 9/9/2024 | 7 |
| 1 | F | Albino | 7/8/2024 | 9/5/2024 | 9/9/2024 | 8 |
| 1 | F | Albino | 7/8/2024 | 9/5/2024 | 9/9/2024 | 9 |
| 3 | F | Albino | 7/8/2024 | 9/5/2024 | 9/9/2024 | 11 |
| 3 | F | Albino | 7/8/2024 | 9/5/2024 | 9/9/2024 | 12 |
| 1 | F | Albino | 7/8/2024 | 9/5/2024 | 9/9/2024 | 15 |
| 2 | F | Albino | 7/8/2024 | 9/5/2024 | 9/10/2024 | 6 |
| 1 | F | Albino | 7/8/2024 | 9/5/2024 | 9/10/2024 | 8 |

**Supplemental Table 2:** Mouse numbers by sex, genotype, and treatment (excluding those used in open field). Mouse numbers separated into genotype, treatment arm, and sex. These numbers do not include mice used in the open field assessment.

| Genotype | Treatment | Sex | # of Mice |
| --- | --- | --- | --- |
| WT | Saline | F | 6 |
| WT | Saline | M | 7 |
| Q112/+ | Saline | F | 6 |
| Q112/+ | Saline | M | 8 |
| WT | mASO | F | 7 |
| WT | mASO | M | 6 |
| Q112/+ | mASO | F | 10 |
| Q112/+ | mASO | M | 6 |
| WT | hASO | F | 4 |
| WT | hASO | M | 11 |
| Q112/+ | hASO | F | 7 |
| Q112/+ | hASO | M | 7 |
| WT | Combo | F | 7 |
| WT | Combo | M | 8 |
| Q112/+ | Combo | F | 7 |
| Q112/+ | Combo | M | 6 |

**Supplemental Table 3:** Mouse numbers used in open field by sex, genotype, and treatment. Mouse numbers separated into genotype, treatment arm, and sex. These numbers only include those mice that underwent open field testing.

| Genotype | Treatment | Sex | # of Mice |
| --- | --- | --- | --- |
| WT | Saline | F | 6 |
| WT | Saline | M | 13 |
| Q112/+ | Saline | F | 9 |
| Q112/+ | Saline | M | 7 |
| WT | hASO | F | 8 |
| WT | hASO | M | 7 |
| Q112/+ | hASO | F | 7 |
| Q112/+ | hASO | M | 12 |

**Supplemental Table 4:** Mouse deaths prior to 63-day endpoint. Mouse deaths separated by genotype and treatment arm. All mice were sacrificed from 58-66 days of age. These mouse numbers represent those mice that died prior to their planned endpoint.

| Genotype | Treatment | # of Mice | # Dead Prior to Endpoint | % Dead Prior to Endpoint |
| --- | --- | --- | --- | --- |
| WT | Saline | 13 | 1 | 8% |
| Q112/+ | Saline | 14 | 3 | 21% |
| WT | mASO | 13 | 3 | 23% |
| Q112/+ | mASO | 16 | 7 | 44% |
| WT | hASO | 15 | 2 | 13% |
| Q112/+ | hASO | 14 | 1 | 7% |
| WT | Combo | 15 | 1 | 7% |
| Q112/+ | Combo | 13 | 1 | 8% |

**Supplemental Table 5:** SHIRPA assessment scoring. Observational phenotypes assessed during SHIRPA testing in longitudinal ASO study. The higher the score in each respective observation indicates a worsening of phenotype.

| Test | Score = 0 | Score = 1 | Score = 2 | Score = 3 | Score = 4 |
| --- | --- | --- | --- | --- | --- |
| Piloerection | None | Coat stands on end |  |  |  |
| Lethargy | Moderate scratch, groom, and movement | Vigorous scratch, groom, and rapid/dart movement | Extremely vigorous scratch, groom, and rapid/dart movement | Slow scratch, groom, and movement | No movement or resting |
| Tremor | None, absent | Mild | Marked |  |  |
| Palpebral closure | Eyes wide open | Eyes half closed | Eyes closed |  |  |
| Touch Escape | Moderate (rapid response to light stroke) | Vigorous (escape response to approach) | Mild (escape response to firm stroke) | No response |  |
| Pelvic Elevation | Normal (3mm elevation) | Elevated (more than 3mm elevation) | Barely touches | Markedly flattened |  |
| Tail elevation | Horizontally extended (Normal) | Dragging | Elevated / Straub Tail |  |  |
| Hindlimb clasping | Absent | Tendency | Clenching on midline | Clasping immediately |  |
| Forelimb clasping | Absent | Tendency | Clasping immediately |  |  |
| Gait | Normal | Fluid but abnormal /ataxic | Limited movement only | Incapacitated |  |
| Respiration rate | Normal | Hyperventilation | Gasping Irregular | Slow, shallow |  |

